# More than just density: the role of egg number, food volume and container dimensions in mediating larval competition in *Drosophila melanogaster*

**DOI:** 10.1101/2023.07.26.550621

**Authors:** Srikant Venkitachalam, V. S. Sajith, Amitabh Joshi

## Abstract

Flies of the genus *Drosophila* have been extensively used as model systems to study competition. Experiments using larval crowding in these species have furthered our understanding of competition ecology, stress-adaptation, density-dependent selection and population dynamics. Historically, larval competition has been induced by crowding the larvae in high density cultures, compared to low-density controls. However, recent studies have shown that two larval cultures having the same total eggs per mL food density, with different absolute quantities of eggs and food, can differ greatly in their density-specific fitness functions. Similarly, populations adapted to two types of cultures achieving the same density through different means, can also evolve different traits. Thus, it is clear that there is more to larval crowding than just eggs/food density, which has until now been the benchmark variable for quantifying larval crowding in *Drosophila* studies. In the current study, we explore the consequences of implementing crowding in different ways, using a three-way factorial experiment with egg number and food volume, cast into different food column heights or cylindrical vials with different diameters. We find that not just the same density, but cultures having the same egg number and food volume combination but experienced in food columns of varying height and diameter can have very different pre-adult survivorship and development times. We further propose a new variable for quantifying larval crowding – effective density, which is the density within the larval feeding band, a volume of food close to the surface, with access to air, wherein a majority of the larvae feed. We show that effective density is a much better predictor of the outcomes of competition than the popularly used total eggs/food density.

## Introduction

Competition can potentially play a large role in shaping the ecology of populations and communities (Elton and Miller 1954; Nicholson 1955), and be a major force driving their evolutionary trajectories (Miller 1967; Wilson 2014). The importance of competition as an impetus for natural selection has been highlighted at least since the time of Darwin (Darwin 1859). Considerable research, both theoretical and experimental, has been done to define and understand competition, as well as its role in ecology and evolution (Sang 1949; Birch 1957; de Wit 1960; Bakker 1961; Miller 1967).

Over the last century, flies of the genus *Drosophila* have become some of the most popular laboratory model systems to experimentally study both intra- and inter-specific competition (Pearl and Parker 1922; Moore 1952; Miller 1964; Ayala 1969; Mueller 1985, 1988; Joshi and Thompson 1995). The holometabolous life cycle of *Drosophila* flies has allowed researchers to partition intra-specific competition for growth-related resources in the larval stage, from the competition for mates in the adult stage (Prasad and Joshi 2003). While studies of adult competition through induced adult crowding have also been conducted (Robertson and Sang 1944; Ohnishi 1976b; Joshi et al. 1998), the present study is focused on the larval phase of competition.

Larval competition in *Drosophila* has had a very rich history of study, and has contributed greatly to the understanding of density-dependent effects, density-dependent selection and the consequent population dynamics. The earliest studies showed that larval competition could be inferred from crowding eggs/larvae at high densities and observing the effects on fitness related traits such as pre-adult survivorship, pre-adult development time and size of eclosing adults (Sang 1949; Bakker 1961; Ohnishi 1976*a*). Several studies additionally investigated the differences in larval competitive ability of various strains of *D. melanogaster* (Bakker 1961, 1969; Gale 1964; Kearsey 1965; Mather and Caligari 1981; de Miranda et al. 1991). Models were also created in an attempt to capture the mechanisms of larval competition (Bakker 1961; De Jong 1976; Nunney 1983; Jansen and Sevenster 1997). Subsequently, selection experiments were carried out to adapt populations to chronic larval crowding, resulting in the evolution of increased larval competitive ability (Mueller 1988; Nagarajan et al. 2016; Sarangi et al. 2016, Venkitachalam et al. 2023).

Researchers have also used the effects of larval crowding as proxies for increased stress at the juvenile phase. They have consequently studied the effects of crowding-adaptation on the correlated evolution of thermal stress resistance (Kapila et al. 2021*a*), as well as the immune system (Kapila et al. 2021*b*). Effects of crowding have also been used to explore yeast availability (Klepsatel et al. 2018), thermal stress (Sørensen and Loeschcke 2001; Henry et al. 2018), and larval crowding has enabled the manipulation of adult body size for sexual selection experiments (Mital et al. 2021). A recent study on transcriptomic consequences of larval crowding was also carried out (Morimoto et al. 2023).

Most studies exploring larval competition or otherwise implementing larval crowding have usually employed a single, or a gradient of, ‘high density’ larval cultures as the experimental regime(s), and compared it to a control ‘low density’ regime. While the low-density cultures were relatively consistent – they involved rearing larvae in high food volumes (5-8 mL, depending on the study) with low starting egg numbers (30-80 eggs, depending on the study), the high-density cultures employed in experiments varied a great deal (reviewed in Henry et al. 2018; Morimoto and Pietras 2020). These cultures varied widely in the number of eggs, the food volumes and the container dimensions used – with the focus being on studying the ‘crowded’ phenotype at an arbitrary high larval density, compared to the ‘uncrowded’ phenotype.

A series of long-term selection experiments over the last four decades, studying density-dependent selection using *Drosophila* populations adapted to larval crowding, ultimately challenged the notion that the consequences of crowding could be predicted based on larval density alone (reviewed in Venkitachalam et al. 2022). Initial studies from the 1980-1990s saw a canonical view emerge for adaptation to larval crowding (Mueller 1997; Joshi et al. 2001; Prasad and Joshi 2003; Mueller et al. 2005*b*; Mueller 2009; Mueller and Barter 2015). This view held that, based on empirical observation, a number of traits would consistently evolve in *Drosophila* populations adapted to larval crowding – such as increased larval competitive ability (Mueller 1988), primarily via increased larval feeding rate (Joshi and Mueller 1988, 1996) or increased metabolic waste tolerance (Shiotsugu et al. 1997; Borash et al. 1998), with a trade off in the form of a reduction in food to biomass conversion efficiency (Mueller 1990; Joshi and Mueller 1996), as compared to low-density control populations. Subsequent studies also carried out such larval crowding adaptation experiments – albeit at a different combination of eggs and food than used earlier (Nagarajan et al. 2016; Sarangi et al. 2016). While the older studies crowded larvae in starting cultures of roughly 1500 eggs in 6 mL food in 6 dram vials, the more recent studies employed cultures with approx. 600 eggs in 1.5 mL food, in wider 8 dram vials. These more recent studies resulted in the evolution of traits very different from what the canonical view predicted. While greater larval competitive ability did evolve, there was no change in feeding rate or waste tolerance. Instead, the crowding-adapted populations evolved shorter development time and greater efficiency of biomass accumulation with respect to time (Nagarajan et al. 2016; Sarangi et al. 2016).

Thus, after three decades of selection studies using larval crowding, it was evident that, while most crowding-adaptation experiments would lead to the evolution of increased larval competitive ability compared to the low-density adapted populations, the specific traits that evolved in different populations could vary widely depending on the details of how exactly the larval crowding was imposed (Sarangi et al. 2016; Sarangi 2018).

These effects of the details of how crowding was imposed were further explored in recent years. One study examined if crowded cultures with the same overall density achieved through different egg numbers and food volumes would differ in their density-specific fitness functions (Sarangi 2018). This was done by crowding larvae of low-density reared populations in two cultures at the same overall density: in either 600 eggs in 1.5 mL food or 1200 eggs in 3 mL of food, in the same kind of vial. Quite surprisingly, the two types of cultures with the same density led to very different patterns of pre-adult survivorship and development time (Sarangi 2018), indicating that density-specific fitness functions of two cultures with the same density achieved via different combinations of egg numbers and food volume could be quite different. A subsequent evolutionary experiment also yielded concordant results. Two sets of populations evolved at the exact same density through different egg and food combinations (600 eggs in 1.5 mL food vs. 1200 eggs in 3 mL food) led to the evolution of differences in terms of feeding rate, pre-adult development time (Sarangi 2018), egg size and egg hatching time (Venkitachalam et al. 2022). These populations also showed differences in the effectiveness and tolerance components of larval competitive ability across a gradient of high-density cultures (Venkitachalam et al. 2023).

The significance of these findings can be understood by applying them to most of the earlier studies carried out on the effects of larval competition and comparisons of competitive ability. How could the results across several studies applying a range of different high-density conditions be comparable, when the same density in closely related populations resulted in large differences in density-specific fitness-functions? Clearly, the commonly used parameter of total density of eggs per unit volume of food itself is not enough to predict the ecology or evolution of traits in larval crowding. There are likely other factors that also influence the ecological and evolutionary outcomes of larval competition beyond the total density of a crowded culture.

The current study aims to explore the aspects of larval crowding beyond the total eggs/food density that may be important in determining the density-specific fitness functions of crowded cultures. For this purpose, we studied two fitness related traits that have been shown to change under the influence of larval crowding – pre-adult survivorship and pre-adult development time (Sang 1949; Bakker 1961; Ohnishi 1976a) – in order to ask a fundamental question: how do these fitness-related traits change with density, when the density change is accomplished by varying different parameters like egg number, food column height and food column diameter?

Some clues to a better understanding of the different facets of larval density are provided by the extensive studies by J. H. Sang in the 1940s. His initial studies showed that keeping all else equal, food volume and egg number could both be varied to change the degrees of larval crowding. While this is now well understood, and has remained the basis of work on larval crowding ever since, Sang also studied two different implementations of changing food volume – firstly, keeping the same cylindrical container and changing the height of the food column (which changes the food volume) and secondly, changing the width (i.e. cross-section surface area) of the cylindrical container itself, while keeping the same food column height. He found that these two ways of implementing crowding led to different outcomes in terms of survivorship, development time and weight of adult flies (Sang 1949). He further calculated that the same food volume cast in different dimensions could have different effects of surface area and food column height (Sang 1949). It should additionally be noted that most subsequent studies altered food volume primarily via changing food column height in an unchanging container type. Unfortunately, these secondary results from Sang’s studies appear to have been ignored due to the importance of the primary result from the research article, which was one of the first demonstrations of the general effects of larval crowding in *D. melanogaster* (Sang 1949).

In the current study, we performed a rigorous follow-up of Sang’s finding on the effects of different ways of implementing crowding. We conducted a fully factorial study of the three factors that can influence larval crowding – 1) egg number, 2) food volume via changing food column height, and 3) food volume via changing the surface area of food exposed to air.

A fully factorial design allowed us to set up several comparisons which within themselves had the same combination of egg number and food volume, but were cast in different cylindrical container dimensions. We asked if there are any differences to be found between any of these cultures. We further explored why there might be reason to find any differences between cultures that essentially appear to be the same implementation of crowding in terms of egg number and food volume, ignoring the container.

## Methods

### Populations used

We used four replicate large (1500+ adults), discrete generation, laboratory *D*. *melanogaster* populations (designated MB_1-4_) whose ancestry and maintenance have previously been described in detail (Sarangi et al. 2016). These populations have also served as ancestral controls to three sets of larval crowding-adapted populations (see Venkitachalam et al. 2022), and are thus maintained at a relatively low larval density. We started each generation of maintenance by collecting eggs from adults of the previous generation in vials at a low density of 60-80 eggs in ∼6 mL of cornmeal-sugar-yeast medium food (see Sarangi et al. 2016) in cylindrical glass vials approx. 95 mm tall, with an inner diameter of around 22.5 mm^2^, resulting in a cross-section surface area of approx. 400 mm^2^ (the food column height was approx. 15-20 mm). After egg collection, we waited until 11 days for the metamorphosis to complete for most of the individuals, and transferred the eclosed adults of each population to a respective Plexiglas cage (25 x 20 x 15 cm^3^). We provided fresh cornmeal medium food plates to the adults in the cages on days 11, 12, 14 and 17 from the day of egg collection (day 1), as well as a moist cotton ball to maintain high relative humidity (>70%). On day 18 from egg collection, we replaced the food plate with a fresh one covered in a yeast-water paste mixed with a drop of glacial acetic acid. On day 20, we removed the yeast-covered plate and provided a food plate cut into two parts for 18 hours. This provided a vertical surface to the *Drosophila* for easier egg laying. Finally, on day 21, we removed the two food plate sections, now heavily laden with eggs. The collection of eggs from these plates served as day 1 for the next generation. This completed a single generation of maintenance. All populations were maintained in constant light (LL), at around 25°C, and a relative humidity of 70-90%.

### Collection of eggs for experiment

For the start of the assay, we singled out one replicate population per generation and collected an additional 80 vials (each with 60-80 eggs, ∼6 mL food, approx. 400 mm^2^ cross section surface area per vial). On day 11 from egg collection, we transferred the eclosed adults from the extra vials to two new cages – adults from 40 vials were transferred to each cage, at random. These cages were each provided a food plate covered with yeast-water paste (containing a drop of glacial acetic acid). On day 14, we provided the cages with fresh cornmeal plates for 1 hour with vertical edges for egg laying. These were discarded after removal from the cages, as they served to collect the eggs that were previously incubated in the females. This would ensure that the subsequent eggs laid by the females would be freshly fertilised, and no inadvertent head starts due to early hatching would be provided to the competing larvae in the experiment. Following plate removal, we provided another egg collection plate to each cage for 4 hours. These were harder plates, containing twice the usual amount of agar (2%), with only yeast and sugar added, and facilitated easier removal of eggs for counting (see Sarangi 2018 and Venkitachalam et al. 2022). After 4 hours, we removed the egg collection plates, and used the eggs laid thereon for the experiment. We added another one of the harder yeast-sugar-agar plate to each cage for 4 hours, in case the first batch did not yield the required number of eggs. The eggs were removed from plates of both cages and were transferred to a non-nutritive 1% agar plate (see Venkitachalam et al. 2022), and thereafter thoroughly mixed prior to starting the egg counting.

After removal of the second harder agar plate, we provided another food plate covered with yeast paste to each cage. On day 15 from egg collection, we removed the yeast plate and repeated the egg collection process of day 14. Eggs taken on day 14 were assigned the Day 1 factor level, and those taken on day 15 were assigned the Day 2 factor level (see below).

### The experimental set up

In the introduction, we listed three ways in which the degree of larval crowding in cylindrical culture vials may be affected, assuming all else is kept unchanged (fig. 1B):

1. Changing starting egg number.
2. Changing food volume via food column height, keeping the same container dimensions.
3. Changing food volume via changing the diameter of the cylindrical vial, thereby changing the surface area of food in contact with air.

**Figure 1.**
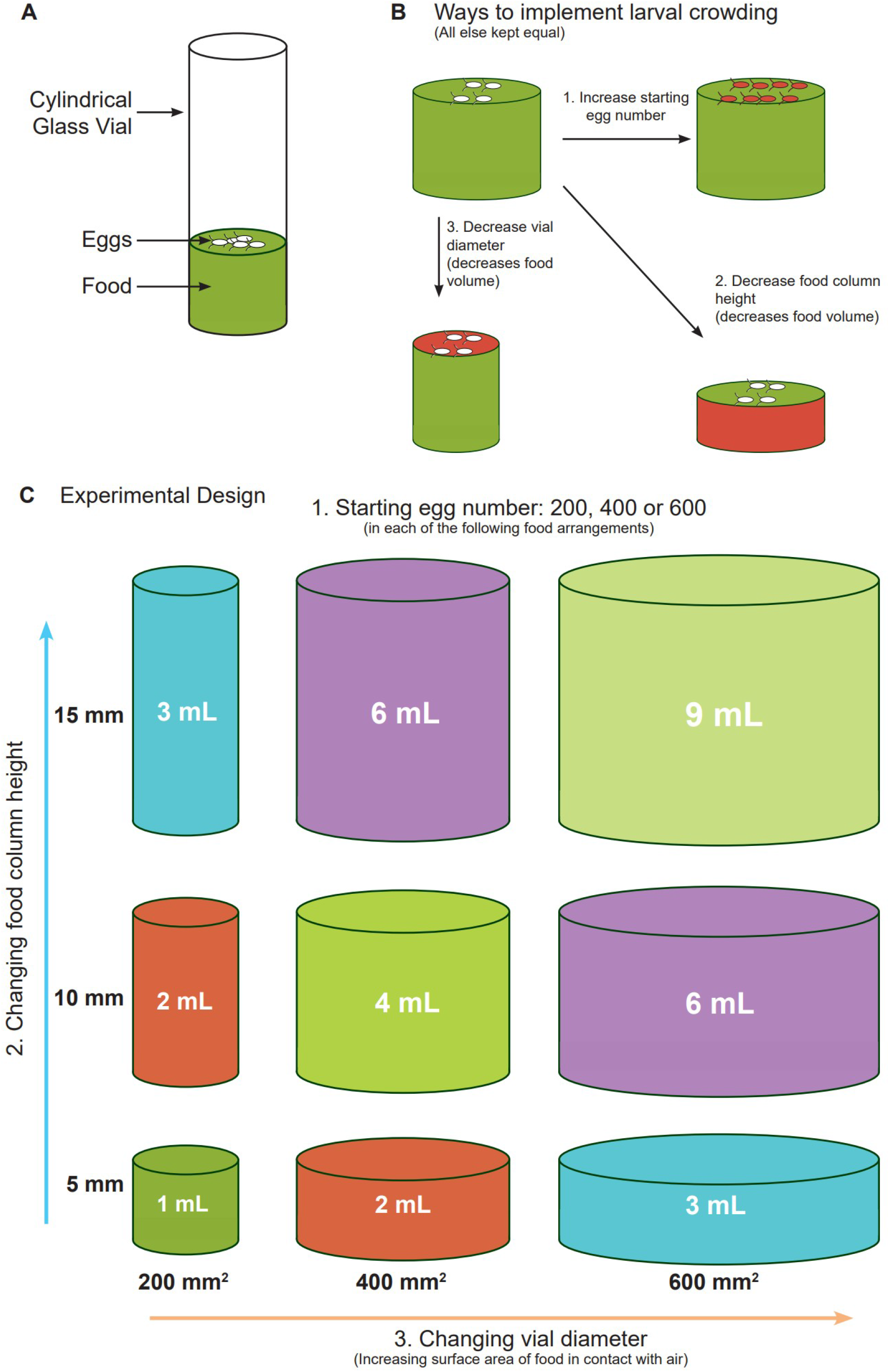
The complete experimental design. A) diagrammatic representation of a cylindrical culture vial. B) The three ways of imposing crowding, assuming all other conditions are constant. C) The design of the experiment, with three levels each of three factors used 1. Egg number; 2. Food column height; 3. Vial cross-section surface area (affected by vial diameter).

We used the collected eggs on 1% agar to conduct an experiment incorporating all three crowding-inducing variables in a three-way, fully factorial design.

1. The eggs were counted exactly for three levels of starting egg numbers: 200, 400 and 600.
2. We used three levels of food column height (∼5 mm, ∼10 mm and ∼15 mm): hereafter referred to as column height.
3. We used three levels of vial cross section surface area (∼200 mm^2^, ∼400 mm^2^ and ∼600 mm^2^): hereafter referred to as surface area.

The complete experimental design is detailed in figure 1. Using three levels of column height and three levels of surface area in a factorial design gave rise to a total of 9 combinations of culture types (fig. 1C). Among these 9 combinations, the food volumes of 2, 3 and 6 mL were present in two types of culture each (fig. 1C, colour matched). The first type had lower surface area and greater column height; the second type had greater surface area and lower food column height. The remaining 3 food volumes – 1, 4 and 9 mL – were unique.

To each of the 9 combinations of surface area and column height, we added 200, 400 or 600 eggs. There were thus 27 different combinations of egg number × surface area × column height. Subsets of these 27 treatments were used to draw relevant comparisons (see Results).

Each of the 27 treatments had four replicate vials for each of the four replicate MB populations, split across two days of egg collection. Thus, we collected two replicate vials per treatment combination on day 14, as well as two replicate vials on day 15.

In total, each replicate population had 108 vials, for a grand total of 432 vials across replicates. A total of 172,800 eggs were used for the entire experiment.

### Collection of eclosing flies

We collected all eclosing flies from each of the 432 experimental vials, split across four replicate populations, in order to measure pre-adult survivorship, as well as pre-adult development time. For this, we checked the assay vials every twelve hours once the first pupae started darkening. These checks were done starting on the 8^th^ day from egg collection, 12 hours apart. Once the first flies started eclosing (usually on the morning of the 9 ^th^ day), we transferred the flies to an empty vial (each of the 108 vials per replicate population had a corresponding transfer vial of similar dimensions). The flies were sacrificed by dousing the transfer vials in liquid nitrogen. Thereafter, they were removed to a clean white surface and counted. The checks were held 12 hours apart until day 12 from egg collection, following which they were instead conducted 24 hours apart. The vials were checked daily until all eclosion ceased.

The total number of adults eclosing from a vial, divided by the number of eggs added to the starting culture, was the measure of pre-adult survivorship per vial. The development time information of each fly was obtained by the total time taken from the mid-point of the parental egg laying interval to the mid-point of the time-window within which that individual eclosed.

### Statistics

We carried out fully factorial mixed model Analysis of Variance (ANOVA) on each of pre-adult survivorship per vial and pre-adult development time (both mean and variance per vial). Survivorship was also arcsine square root transformed and analysed to verify the results seen with non-transformed values.

All ANOVA were carried out on Statistica for Windows (Statsoft 1995). Egg number, food column height and vial cross-section surface area were treated as fixed factors with three levels each in the analysis, respectively. Additionally, day of experiment set up (day 14 or day 15 from egg collection) was also a fixed factor with two levels. Finally, population replicate number was treated as a random block factor with four levels. Pairwise post-hoc comparisons were done using Tukey’s HSD.

We also carried out linear regression of the pooled data on pre-adult survivorship, mean pre-adult development time, variance of pre-adult development time and coefficient of variation of pre-adult development time, respectively. These were each treated as response variables against either total density or effective density as predictor variables (see results section).

The plotting of data was done on R release 4.2.2 (R Core Team 2022), using ggplot2 and tidyverse packages (Wickham 2016; Wickham et al. 2019).

## Results

### Main effects

Each of the three factors of interest – egg number, column height and surface area had a significant main effect on pre-adult survivorship, as revealed by the ANOVA (table s1). On an average, increasing the number of eggs, as well as decreasing the volume of food through reduction in either surface area or column height decreased pre-adult survivorship (fig. s1a-c). Although all three effects were significant for survivorship (table s1), the magnitude of change of each outcome with the factors was different. While large changes in survivorship were seen across the three levels employed for egg number (fig. s1a) and surface area (fig. s1b), column height induced relatively smaller changes in survivorship with changing levels (fig. s1c).

The pattern for mean pre-adult development time differed based on the factor tested, although the ANOVA revealed a significant main effect for each factor (table s2). In case of egg number, mean pre-adult development time decreased with decreasing egg number (fig. s1d). For surface area, decreasing levels resulted in an increase in mean development time (fig. s1e). However, for column height, decreasing levels saw a small decrease in mean development time (fig. s1f). Thus, averaged over all other factors, an increase in crowding through reduced column height resulted in a slightly shorter development time. It should, however, be noted that the extent of differences in mean development time induced by the changing column height were relatively minor as compared to those induced by changes in both egg number and surface area. Variance of development time showed similar patterns as the mean (fig. s1g-i); however, there was no significant main effect of food column height (fig. s1i)

### Comparisons of cultures with the same egg number and food volume in containers of different diameters

Different patterns of development time via increasing surface area vs. column height indicated some fundamental difference in how larval crowding was experienced as a result of reduced food volume achieved by these two modes. This was further explored by looking at subsets of comparisons for the three-way interaction of egg number × column height × surface area. These interaction effects were statistically significant for all three outcomes of crowding investigated (tables s1, s2, s3).

Subsets of comparisons were made among cultures with the same food volume and egg number, but across different vial diameters. As seen in figure 1C, the food volumes of 2, 3 and 6 mL are present in two culture types – one with larger surface area and reduced column height, and the other with smaller surface area and a greater column height. Thus, for each of the three egg numbers used – 200, 400 and 600 – there are two cultures at 2 mL, 3 mL and 6 mL respectively, with not just the same overall density, but the exact same combination of egg number and food volume, achieved via different combinations of surface area and column height.

Figure 2 shows the mean pre-adult survivorship and mean pre-adult development time in the contrasting cultures of both 2 mL (fig. 2A and 2C) and 3 mL (fig. 2B and 2D) across increasing egg numbers. In case of the 2 mL cultures, when increasing the starting egg numbers, the culture with the narrow surface area and greater height (coloured blue) showed a greater loss in survivorship (fig. 2A) and a greater increase in mean development time (fig. 2C), compared to the culture with the greater surface area and shorter column height (coloured orange). These differences between the ‘blue’ and ‘orange’ cultures were further exacerbated at the 3 mL food volume (fig. 2B and 2D).

**Figure 2.**
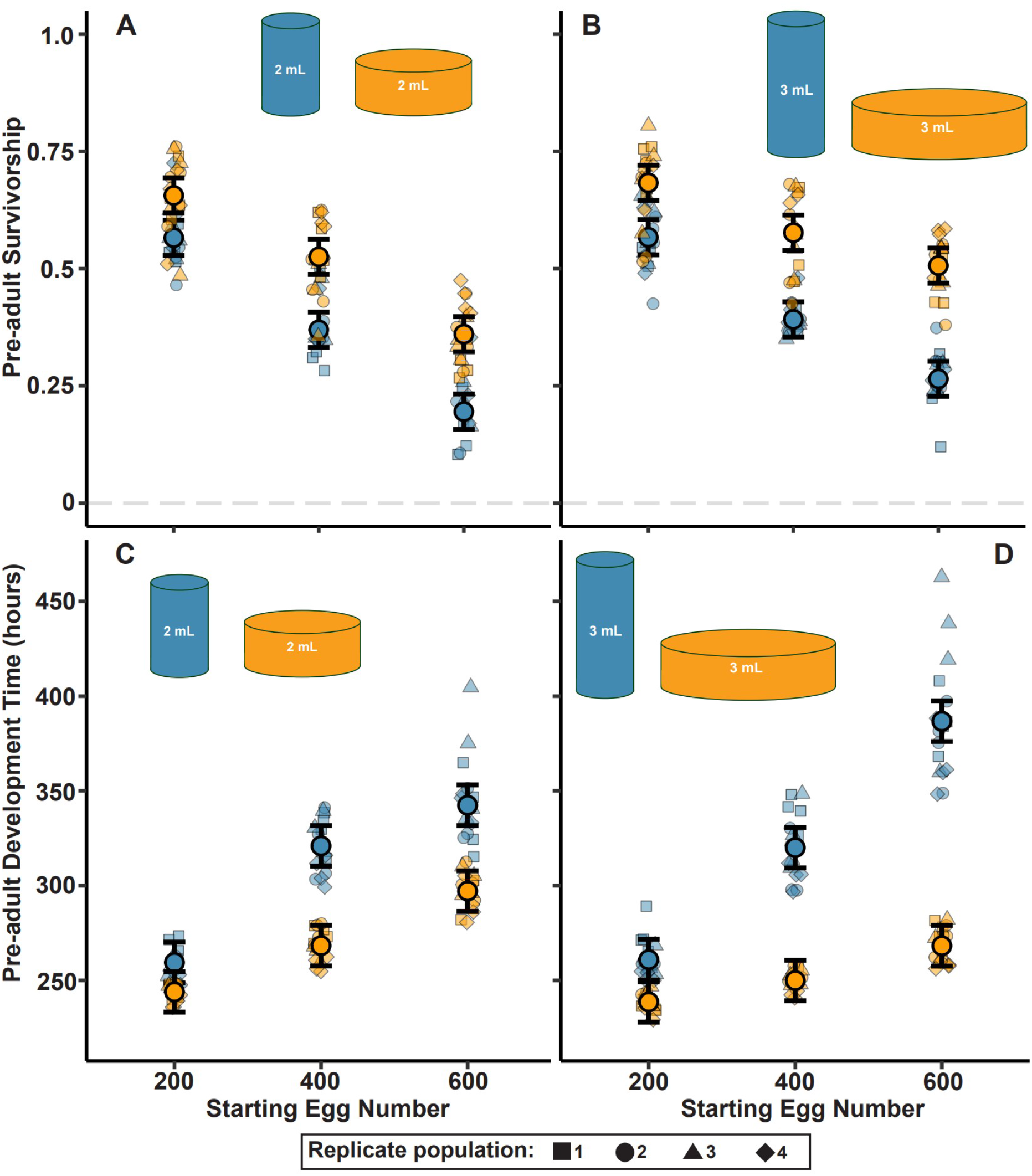
Mean pre-adult survivorship and mean development time at the same food volume. The food volumes and shapes into which they are cast are subsets of the experiment design showed in fig. 1C. In the current figure, A, C denote 2 mL; B, D denote 3 mL starting food volume. The error bars denote 95% C.I. around the overall means for the respective traits at each egg number, for the Tukey’s post-hoc test.

Similar results were also seen in the case of 6 mL food, although the differences were relatively smaller in magnitude, and often non-significant (fig. s2a, s2c). The variance in development time also showed differences, very similar in pattern to the average development time, in most cases (fig. s3a, s3b, s3). Additionally, we have also plotted the data for 1 mL, 4 mL and 9 mL cultures for survivorship (fig. s2b), as well as the mean and variance in development time (fig. s2d, s3d).

### Larval feeding band, total density and effective density

Our results quite clearly showed a large difference in survivorship and development time – and hence the underlying fitness functions – of two cultures that had the same egg number and food volume, with the main differentiating factor likely to be the surface area and column height that the food was cast into. To further investigate the possible role of surface area and column height, we focused on the larval ‘feeding band’ in highly crowded cultures. We define the feeding band as the volume of food close to the surface, with access to air, to which larval feeding is usually restricted (fig. 3A), implying that the maximal depth of food that a larva can access is roughly equal to the length of the larva. Consequently, relatively shallow food columns should allow the larvae to access most of the food (fig. 3Aiii), whereas, high food columns that extend beyond the feeding band are likely to have some food at the bottom that would not be accessible by the larvae until the top layer of starting food is sufficiently depleted (fig. 3Ai). Thus, two cultures with the same food volume that differ in surface area and food column height, may have very different volumes of initially accessible vs. inaccessible food, assuming similar feeding band depths. This is likely what we see in our experimental design, with food volumes of the 2, 3 and 6 mL combinations (fig. 2, s2). For example, in the 3 mL culture contained within the relatively narrow vial (∼200 mm^2^ surface area), the food column was relatively high (∼15 mm) (fig. 2B, 2D; ‘blue’ cultures). In contrast, for the 3 mL culture contained within the wider vial (∼600 mm^2^ surface area), the food column was much shorter (∼5 mm) (fig. 2B, 2D; ‘orange’ cultures). Given the shallow depth of food in the wider vial, larvae could possibly access a lot more of the food at any given time, compared to larvae in the narrow vial which might not be able to penetrate to the depths of 15 mm.

**Figure 3.**
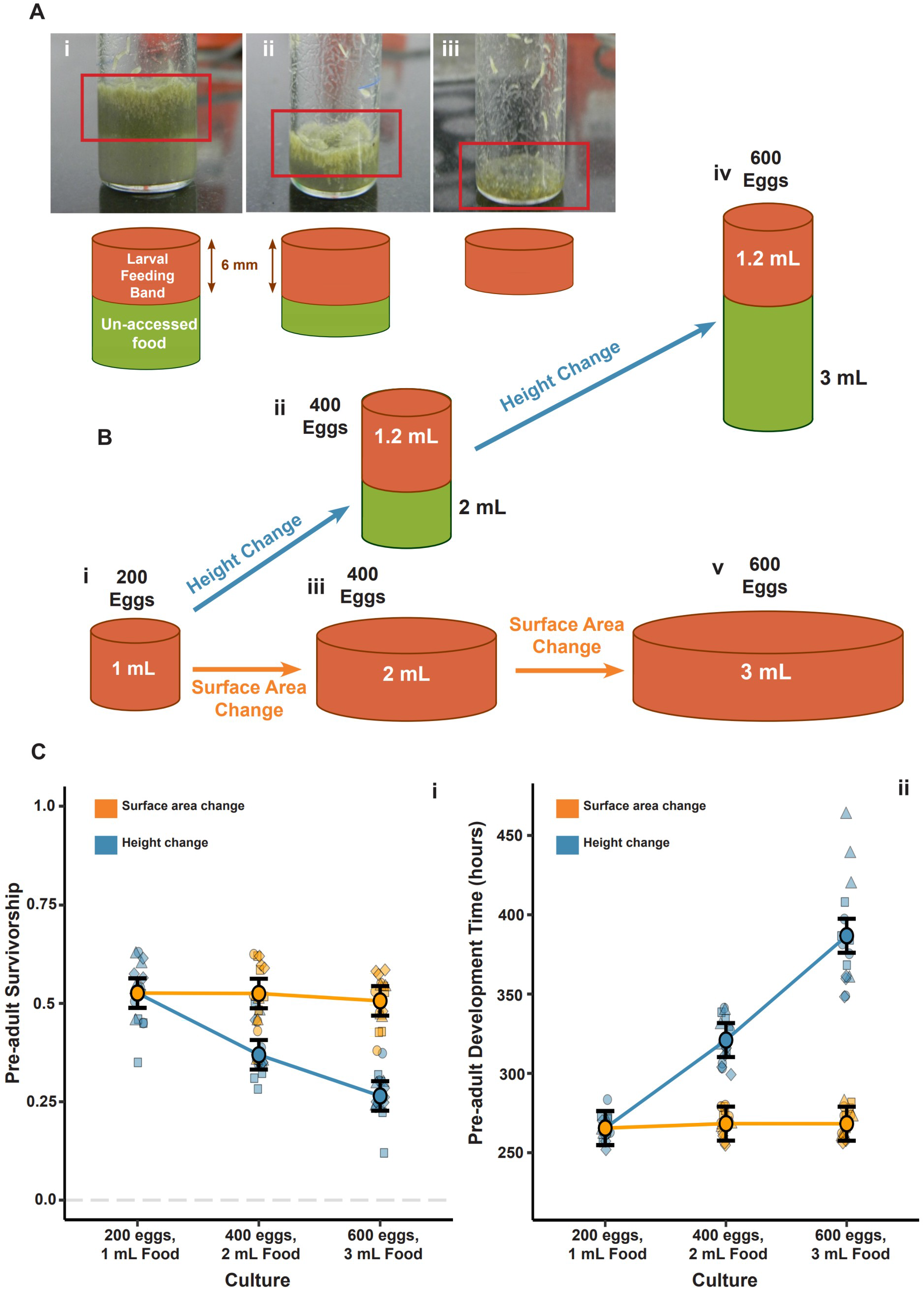
A. The presence of the feeding band and its diagrammatic representation across three food column heights. In case of tall food columns (i and ii), the food at the bottom in not accessed by larvae until the top layer of food is depleted. In shallow food columns (iii), the entire food volume comprises the feeding band. B. For a total eggs/food density of 200 eggs/mL, there are 5 ways to achieve it by altering egg and food quantities, in our experimental design (i-v), as shown in the diagram. If food column height is increased (ii, iv), the feeding band is restricted to 1.2 mL food, thus changing the effective density in the feeding band (fig. s4). In case surface area is increased (iii, v), keeping a shallow food column, the total density and effective density are the same (fig. s4). C. The outcomes of crowding at a total density of 200 eggs/mL in 5 different ways as listed in B. (i) shows pre-adult survivorship; (ii) shows pre-adult development time (hours). The orange line represents changing surface area, keeping total density and effective density the same. The blue line represents changing food column height, which results in effective density differing from total density. Error bars represent 95% C.I. around the overall mean for a given culture and vial type, for each respective trait, as calculated from Tukey’s post-hoc test.

For the following distinction between total vs. effective density, we assumed the feeding band depth to be a constant 6 mm, with the caveat that the actual feeding band depth is likely to be a more complex parameter (see Discussion). In addition to the total density (eggs per unit food volume in the vial), we define a new term – effective density – calculated as the number of eggs per unit volume of the feeding band. The effective density better reflects the actual crowding experienced by feeding larvae in the upper 6 mm of the food column.

We then calculated the total and effective density per culture combination for each of the 27 cultures, as shown in fig. s4. Cultures with short food column height, which have the feeding band volume equal to the total food volume end up having the same total and effective densities. Cultures with longer food column heights have feeding band volume as a subsection of the total food (up to 6 mm deep), and thus have higher effective density than total density. For the 3 mL food example, the wider vial has a feeding band that provides access to the entirety of the food (fig. 3Bv), whereas the 3 mL food in the narrow vial has only ∼1.2 mL food accessible in the feeding band (fig. 3Biv), thereby leading to a far higher effective density for each given egg number (fig. s4).

Additionally, the calculated total and effective density for each treatment combination yielded multiple cultures which had the same total density, but different effective densities. Three levels of total density had at least 5 different representatives each. These were: 200 eggs/mL food; 100 eggs/mL food and 66.67 eggs/mL food (fig. 3B, s5A, s5B, colour matched). For example, the first case, a total density of 200 eggs/mL food, could be obtained in the following scenarios:

1. 200 eggs in 1 mL food (fig. 3Bi).
2. 400 eggs in 2 mL food in ∼200 mm^2^ surface area, 10 mm column height (fig. 3Bii).
3. 400 eggs in 2 mL food in ∼400 mm^2^ surface area, 5 mm column height (fig. 3Biii).
4. 600 eggs in 3 mL food in ∼200 mm^2^ surface area, 15 mm column height (fig. 3Biv).
5. 600 eggs in 3 mL food in ∼600 mm^2^ surface area, 5 mm column height (fig. 3Bv).

Similar comparisons for the cases with 100 eggs/mL and 66.67 eggs/mL can be found in supp. figure s5A, s5B, respectively.

The results show that cultures wherein the total and effective density were the same (1, 3, 5 from list above), had very similar pre-adult survivorship as well as both mean and variance of pre-adult development time (fig. 3C, s6). When effective density was different from total density, there were large differences in the pre-adult survivorship, and also in the mean and variance of pre-adult development time (fig. 3C, s6). Similar results were seen for the other total densities as well (fig s5A, s5B).

### Linear regressions: total vs. effective density

To further examine the role of total vs. effective density in mediating competitive outcomes, we also performed linear least-squares regressions by pooling all treatments, with either total density or effective density as the predictor variable. The response variables were either pre-adult survivorship, mean development time per vial, or variance/coefficient of variation in development time per vial. The results can be seen from the table in fig. 4, wherein the *R*^2^ values for both regressions are shown for each response variable. The fitted lines are plotted in fig. s7.

**Figure 4.**
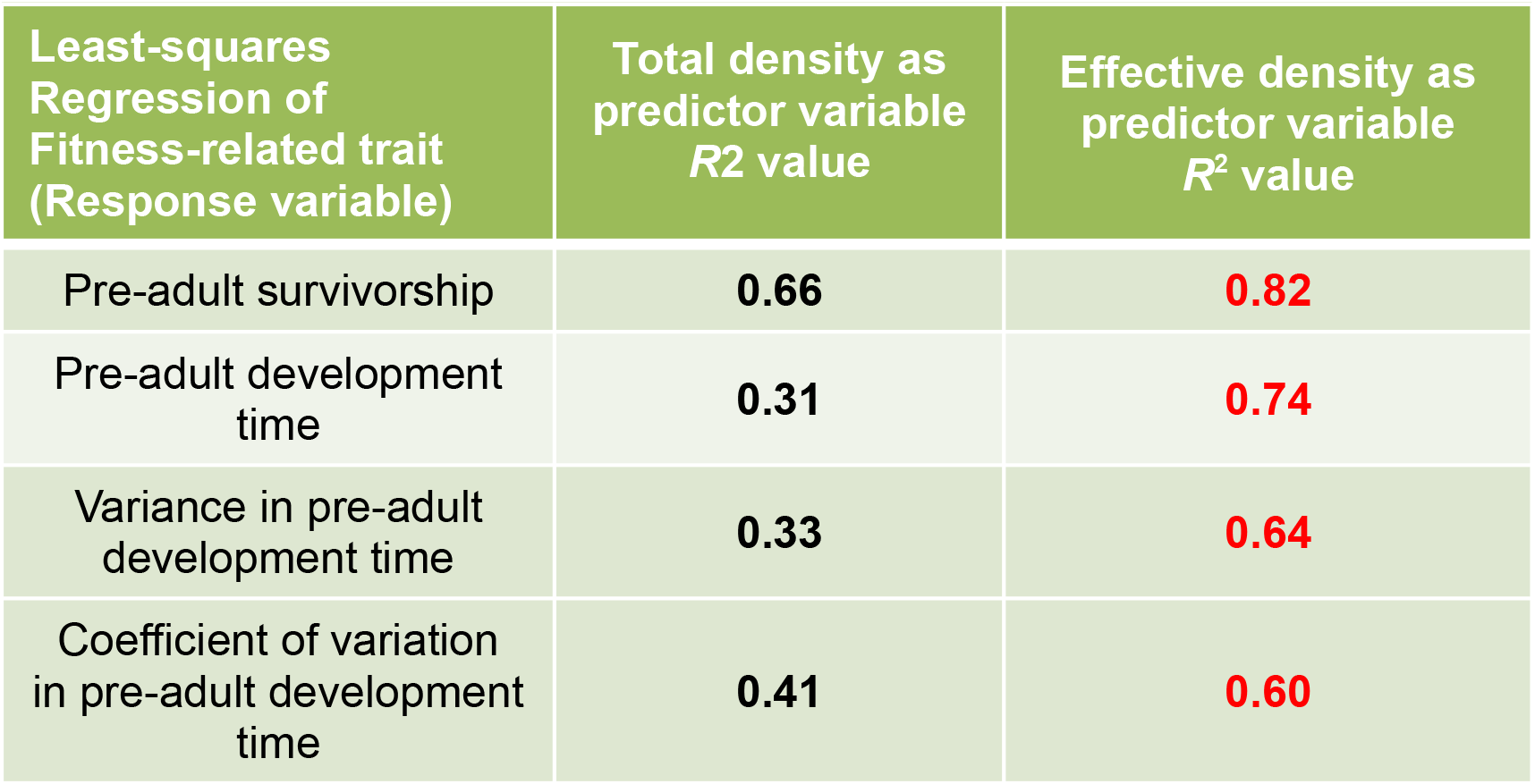
Goodness-of-fit (*R*^2^) values for linear regressions performed by pooling the entire experimental data for each of the outcomes of larval crowding studied as response variables. The predictor variables are either total eggs/food density (left column), or effective (larval feeding band) density.

For each of the traits examined, a linear model using effective density substantially better predicted the response variable than total density. This showed that simple linear models with effective density were better predictors of the outcomes of competition.

## Discussion

We conducted a three-way, fully-factorial experiment on implementing larval crowding in different ways and found differences in density-specific fitness functions of cultures with the same egg and food combination, in different cylindrical dimensions. Across nine pairwise comparisons of the same combination of eggs and food volume, 7 out of 9 showed differences in pre-adult survivorship (fig. 2A, 2B, s2a), and 5 out of 9 comparisons showed differences in mean pre-adult development time (fig. 2C, 2D, s2c). In each case where the comparison yielded a significant difference (and some in which there was a non-significant difference), the culture which had a greater surface area of food in contact with air, with a reduced food column height, had greater survivorship than the same culture cast in a relatively lesser surface area and a greater food column height (fig. 2A, 2B, s2a). Similarly, the cultures with greater surface area and reduced column height also had shorter mean development time, where significant, than their respective counterparts with lower surface area and greater column height (fig 2C, 2D, s2c). These results are also in agreement with the study by Sang (1949). Variance in development time largely followed the same pattern as mean development time (fig. s3a-c).

In order to find out why there were such stark differences simply by property of culture dimensions, we looked at the larval feeding band, assuming a depth of 6 mm as a simplified representative of the depth to which larvae could potentially feed at all times. We thus calculated effective density (initial number of eggs/feeding band volume) for each of the 27 culture types (fig. 1C, s4), in addition to the total density (number of eggs/total food volume), for each culture. We found that while up to five types of cultures could have the same total density by virtue of their eggs and food combinations, their survivorship and development time outcomes were only similar if the effective density was equal to the total density (fig. 3, s5, s6). These strikingly similar outcomes were achieved via an increase in surface area, which kept the total and effective density the same. Additionally, the linear regression of every outcome of larval crowding examined yielded a higher *R*^2^ value when effective density was the predictor variable, as opposed to total density (fig. 4). Overall, these results indicate that effective density plays a far more important role in shaping the outcome of larval competition than total density, despite the latter having been the benchmark for larval crowding till date.

**Figure 5.**
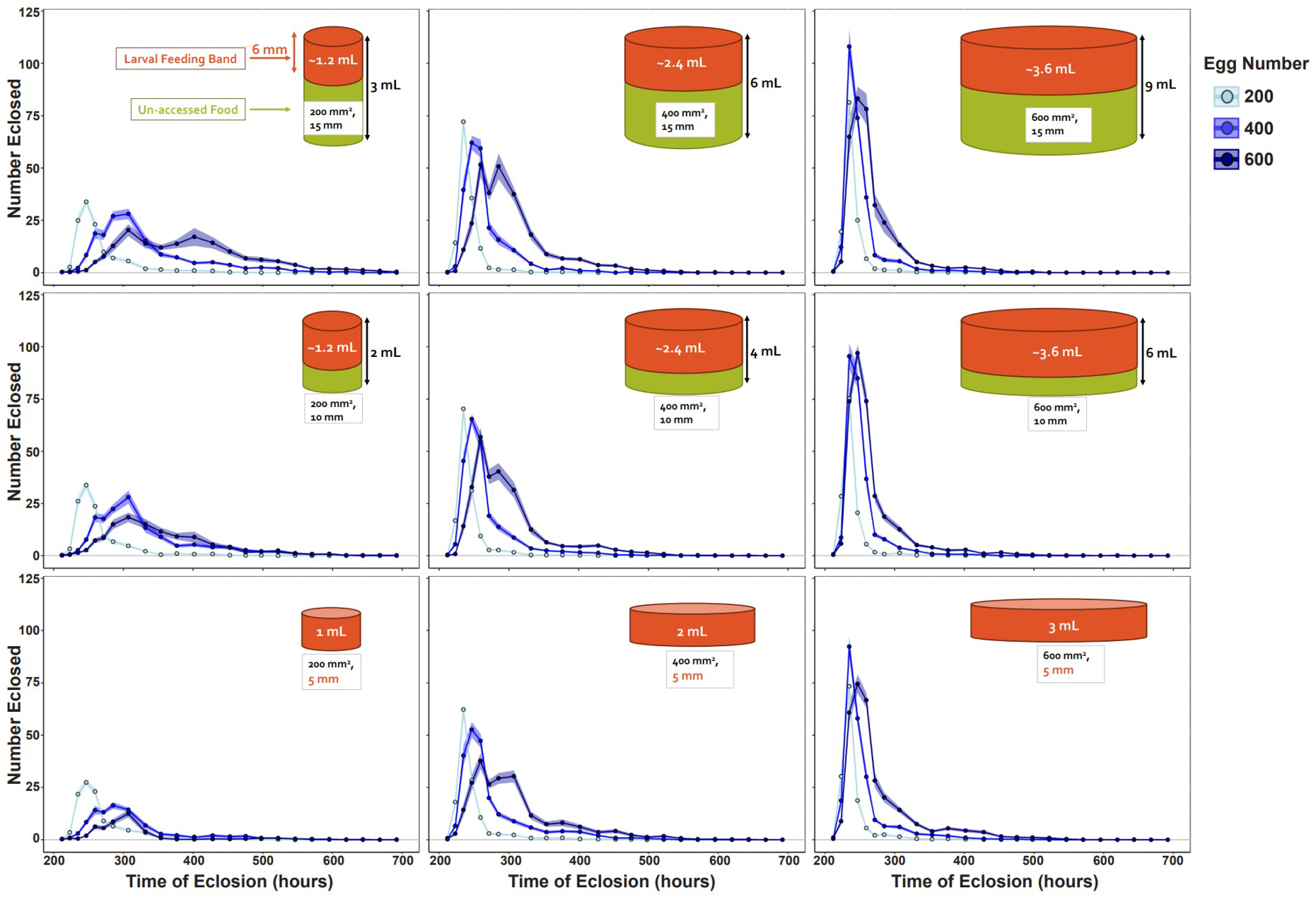
Distributions of time of eclosion of adult flies for each of the 27 treatments used in the current study. The representative culture diagrams are as shown in fig. 1C, with derived feeding band volumes as shown in fig. s5.

Even in terms of the eclosion profiles, effective density seemed to be very important. The eclosion distribution increases in variance at high effective densities coupled with long food columns (fig. 5, s6). At shallow depth of food, eclosion distributions are generally less spread out (fig. 5).

Our results also indicate that beyond effective density, the overall volume and depth of the food column may also help determine the ultimate fate of a crowded larval culture. This is apparent in the comparison of 600 eggs in 2 mL vs. 600 eggs in 3 mL food, cast into 10 mm and 15 mm column heights, respectively (200 mm^2^ surface area each). While both cultures have the same calculated effective density of around 500 eggs per mL of feeding band (fig. s4), the same effective density in a 15 mm food column (3 mL total food) has significantly higher development time (fig. 2) and development time variance (fig s3a, s3b), and a higher, though non-significantly so, survivorship (fig. 2), than in a 10 mm food column (2 mL total food). This does not occur in any other scenario wherein the effective density is the same between two cultures. Thus, for two cultures having the same (high) effective densities, a greater volume of food – achieved via increased food column height – may lead to an average delay in development time along with an increase in variance, perhaps at the benefit of a slight advantage in survivorship. This needs to be explored further, with greater effective densities tested at several food levels beyond those used in the current study.

Due to logistical limitations, the measurement of a 6 mm deep feeding band is a necessary simplification of a complex phenomenon. An early study on larval length of *D. melanogaster* ascertained the maximum measurement to be around 4.5-4.6 mm, which is well within the 6 mm cut-off (Alpatov 1929). We also note that in his studies, Sang (1949) had discussed the existence of the feeding band as well, defining it at 5 mm deep. Given that 5-6 mm is the longest depth to which maximally grown larvae can extend (notwithstanding digging), we assumed 6 mm to be the liberal estimate of the feeding band depth. At the start of the culture, the freshly hatched larvae would be expected to feed in a much shallower feeding band than 6 mm. The feeding band would increase as the larvae grow and gain longer lengths and greater reach. Thus, assuming a depth of 6 mm as constant, any differences captured at this simple calculation are likely to be exacerbated with more precise measurements. Moreover, such precise measurements may be difficult to carry out – realistically, the larval feeding band is likely to be complex over space and time. Spatially, the larvae tend to cluster themselves into feeding grooves near the surface, rather than spread themselves out uniformly over the feeding band (previously observed by Gilpin 1974, Chippindale et al. 1994, Mueller and Barter 2015, S. Venkitachalam, personal observation; also see Gregg 1990). Variable digging length may also be a possibility, although in overcrowded feeding clusters, the larvae could face the risk of drowning due to spatial constraints. The effective larval density of a culture would itself be dynamic and changing throughout the life of the culture, as the larvae grow, drown or leave the culture. This would make the prospect of exactly calculating the feeding band depth or the effective density very difficult.

Given that this study has formally added a new descriptor of larval crowding, it is important to apply this insight to any future studies that aim to implement any form of high density to larval cultures. It becomes extremely relevant to list out not just the number of starting individuals and the total food volume (as well as the type of food used), but also the details of the container – the best practice would be to list the effective density or the average food column height and the surface area of food with access to air. Even in case of studies in nature (Atkinson 1979; Morimoto and Pietras 2020), this finding is likely relevant, as the density of larvae in a fruit is likely to vary across its dimensions, creating local zones of high and low effective densities within a single fruit itself.

Future studies on larval crowding in *D*. *melanogaster* across research labs worldwide should ultimately test the inferential reach of our findings, although we suspect that major exceptions to our results will be rare (and interesting). Besides J.H. Sang, who found similar results in very different strains, another study made note of the effects of surface area on larval competition (Scheiring et al. 1984). Several other studies discussed the possibility of the effects of surface area on the outcomes of larval crowding (Bakker 1961; Gilpin et al. 1976; Chippindale et al. 1994). The literature on larval crowding is also populated with references to un-accessed food at the bottom of high-density cultures with relatively high food volumes (Gilpin 1974; Bierbaum et al. 1989; Mueller 1990).

In earlier studies, *D. ananassae* and *D. nasuta* populations showed similar adaptations to larval crowding in a shallow depth of food with low food volume as *D. melanogaster* (Nagarajan et al. 2016, Sarangi et al. 2016). It remains to be seen whether they would follow similar patterns with respect to the predictability of crowding outcomes by effective density, as seen in the current study. Finally, there have also been findings of the effects of surface area in larval systems from other holometabolous insect clades, such as the moth species *Ephestia cautella* (Smith 1969), *E. kuehniella*, *E. elutella* and *PIodia interpunctella* (Bell 1976), as well as the flour beetles *Tribolium castaneum* (Wool 1969). These studies indicate that feeding-band-like features may exist across several insect groups, although the exact consequences of such a feature may depend on the biology of the species being crowded. However, this sets up the background for potentially interesting future studies that can explore the complex aspects of larval competition and crowding across species.

Mechanistically, it is likely that constrained space in the feeding band at high effective densities may be one major factor driving the outcome of *Drosophila* larval competition. In high food columns with small surface area, most larvae may be unable to access food beyond the depth of the feeding band. Moreover, a large number of larvae trying to feed in a constrained space likely leads to space limitation, despite potentially large volumes of food remaining uneaten at the bottom of the food column. On the other hand, high food columns may have the possible benefit of the diffusing of harmful metabolic waste products away from the feeding band and into the depths of the yet-inaccessible food (Sarangi 2018). This may explain the large variance in egg to adult development time in 600 eggs at 3 mL food at the 15 mm food column length (fig. 3Civ). Under such high effective densities, some space constrained larvae may survive on low amounts of food until most other competitors have left the food, drowned or been poisoned due to high waste levels. Thereafter, these surviving larvae could resume feeding (provided they can tolerate the elevated waste levels), leading to an extended developmental period. Alternatively, some larvae may feed faster and dig deeper into the un-accessed food, leading to a faster development time. Both may be viable strategies, but while the former scenario would delay developmental time, the latter scenario would select for faster digging, feeding and growth. This would ultimately lead to a culture with both increased mean and variance in development time. Our reasoning is also in agreement with some previous studies, which implemented larval crowding at approx. 1500 eggs in 6 mL food cast in high food columns with relatively narrow cross section surface area (Borash et al. 1998; Mueller et al. 2005*a*; Mueller and Barter 2015). Thus, these studies inadvertently used high effective density cultures with long food columns, achieving very large variance in development time. The authors found that early eclosing flies had offspring with increased feeding rate while late eclosing flies had high waste tolerance, as compared to the individuals of the non-selected control populations (Borash et al. 1998). In contrast, in high effective density cultures with shallow food column heights, the food would run out much faster, due to a large number of larvae accessing all of a limited quantity of food. Space constraints might be secondary to food constraints, leading to a relatively truncated feeding period, thus reducing both the mean and variance of the pre-adult development time. This is likely what was observed in later selection studies on chronic larval crowding in *D. ananassae*, *D. nasuta* and *D. melanogaster* (Nagarajan et al. 2016, Sarangi et al. 2016). Most crowded cultures used in laboratories, and perhaps even in nature, likely lie between these two extreme conditions, often leading to a lack of consensus on the effects of crowding and competition.

Several avenues can be explored for future studies. With the experimental design used in the current study, we can also potentially obtain dry weight data for freshly eclosed adults, and map dry weight distribution over time for each culture type. Both mean and variance of dry weight per vial can be measured, as was done for pre-adult development time in the current study. The body weight of emerging flies is one of the most sensitive indicators of the existence of competition (Sang 1949; Bakker 1961; Santos et al. 1997). Moreover, studies have indicated that elongated development time distributions can give rise to differences in adult weight depending on the time of eclosion (Hughes 1980; Sarangi 2018, Venkitachalam et al. 2023). This would be especially relevant for comparisons between cultures which have widely varying development time distributions (fig. 5).

We also know that populations adapted to high effective density cultures in shallow food column depths (total density similar to effective density) differ in evolutionary trajectory from populations adapted to high effective density in longer food columns (total density differing from the effective density) (Sarangi et al. 2016, Sarangi 2018, Venkitachalam et al. 2022). Given the framework we have established, we can test whether these two kinds of population sets differ in their competitive ability depending on the type of culture used. A recent competitive ability experiment carried out by us tested each of the different crowding adapted population sets in varying volumes of food (with shallow food column depths), and the results indicated the existence of culture-dependent competitive ability (Venkitachalam et al. 2023).

Finally, starting new populations within this factorial framework would allow us to adapt populations to very different culture ecologies while keeping the same egg number and food volumes. While logistically intensive, this would be the best test for whether different crowding ecologies can lead to specialised competitive ability i.e., populations exhibiting higher competitive ability under the precise conditions in which they experienced crowding. It may be possible that greater digging distance may be important in competing in high effective density cultures with higher food columns. However, a previous study did not find differences in average digging length in populations adapted to crowding under relatively large volumes of food (Mueller 1990).

In conclusion, we have described effective density as the density of larvae in the feeding band – a volume of food close to the surface of food with access to air, where the larvae feed. The effective density is a far better predictor of three fitness-related outcomes of larval crowding compared to the total density, which is the total number of eggs in the total volume of food of a culture. This finding has important implications for the design of future studies on larval competition in *Drosophila* and perhaps other clades as well.

## Acknowledgements

We thank Profs. T.N.C. Vidya (JNCASR) and Sutirth Dey (IISER Pune) for helpful discussions on the results, and Rajanna N. and Muniraju for help with the experiment. S. Venkitachalam was supported by a doctoral fellowship from JNCASR. This work was supported by a J. C. Bose National Fellowship from the Science and Engineering Research Board, Government of India, to A. Joshi, a Department of Biotechnology, Government of India - JNCASR project, “Life Science Research, Education & Training”, and, in part, by A. Joshi’s personal funds.

## Appendix Supplementary Material

**Table s1.** ANOVA results for mean pre-adult survivorship.

Appendix: Supplementary Material
| Effect | *df* | MS | *F* | *P* |
| --- | --- | --- | --- | --- |
| Eggs | 2 | 2.5593 | 400.881 | < 0.0001*** |
| Surface Area | 2 | 2.5436 | 222.149 | < 0.0001*** |
| Food Height | 2 | 0.2061 | 62.624 | < 0.0001*** |
| Day | 1 | 0.3686 | 6.740 | 0.0806 |
| Surface Area × Food Height | 4 | 0.0357 | 25.669 | < 0.0001*** |
| Surface Area × Day | 2 | 0.0878 | 16.405 | 0.0037** |
| Food Height × Day | 2 | 0.0045 | 0.824 | 0.4828 |
| Surface Area × Eggs | 4 | 0.1565 | 32.952 | < 0.0001*** |
| Food Height × Eggs | 4 | 0.0197 | 6.552 | 0.0049** |
| Day × Eggs | 2 | 0.0682 | 23.124 | 0.0015** |
| Surface Area × Food Height × Day | 4 | 0.0004 | 0.166 | 0.9515 |
| Surface Area × Food Height × Eggs | 8 | 0.0090 | 3.425 | 0.0091** |
| Surface Area × Day × Eggs | 4 | 0.0039 | 2.542 | 0.0944 |
| Food Height × Day × Eggs | 4 | 0.0029 | 1.264 | 0.3371 |
| Surface Area × Food Height × Day × Eggs | 8 | 0.0037 | 1.623 | 0.1704 |

**Table s2.** ANOVA results for mean pre-adult development time.

| Effect | *df* | MS | *F* | *P* |
| --- | --- | --- | --- | --- |
| Eggs | 2 | 101652 | 351.39 | <0.0001*** |
| Surface Area | 2 | 125726 | 468.79 | <0.0001*** |
| Food Height | 2 | 2356 | 6.79 | 0.0288* |
| Day | 1 | 1 | 0.00 | 0.9488 |
| Surface Area*Food Height | 4 | 4716 | 19.56 | <0.0001*** |
| Surface Area*Day | 2 | 3 | 0.01 | 0.9868 |
| Food Height*Day | 2 | 0 | 0.00 | 0.9983 |
| Surface Area*Eggs | 4 | 11565 | 48.32 | <0.0001*** |
| Food Height*Eggs | 4 | 2284 | 16.48 | <0.0001*** |
| Day*Eggs | 2 | 70 | 0.90 | 0.4553 |
| Surface Area*Food Height*Day | 4 | 81 | 0.83 | 0.5330 |
| Surface Area*Food Height*Eggs | 8 | 2831 | 13.27 | <0.0001*** |
| Surface Area*Day*Eggs | 4 | 74 | 0.37 | 0.8276 |
| Food Height*Day*Eggs | 4 | 137 | 1.23 | 0.3488 |
| Surface Area*Food Height*Day*Eggs | 8 | 165 | 1.34 | 0.2741 |

**Table s3.** ANOVA results for variance in pre-adult development time.

| Effect | *df* | MS | *F* | *P* |
| --- | --- | --- | --- | --- |
| Eggs | 2 | 282326582.8 | 367.422 | <0.0001*** |
| Surface Area | 2 | 404285765.8 | 245.276 | <0.0001*** |
| Food Height | 2 | 2363303.6 | 1.703 | 0.2595 |
| Day | 1 | 1116630.7 | 0.792 | 0.4389 |
| Surface Area*Food Height | 4 | 15152499.2 | 8.261 | 0.0019** |
| Surface Area*Day | 2 | 87741.1 | 0.058 | 0.9441 |
| Food Height*Day | 2 | 1652457.8 | 1.041 | 0.4091 |
| Surface Area*Eggs | 4 | 42424753.5 | 25.181 | <0.0001*** |
| Food Height*Eggs | 4 | 3231437.7 | 2.653 | 0.0852 |
| Day*Eggs | 2 | 2205768.4 | 4.350 | 0.068 |
| Surface Area*Food Height*Day | 4 | 850900.5 | 0.296 | 0.8751 |
| Surface Area*Food Height*Eggs | 8 | 10830484.9 | 4.206 | 0.0029** |
| Surface Area*Day*Eggs | 4 | 684030.1 | 0.828 | 0.5326 |
| Food Height*Day*Eggs | 4 | 2843090.7 | 1.294 | 0.3266 |
| Surface Area*Food Height*Day*Eggs | 8 | 1979722.5 | 1.202 | 0.339 |

**Fig. s1.**
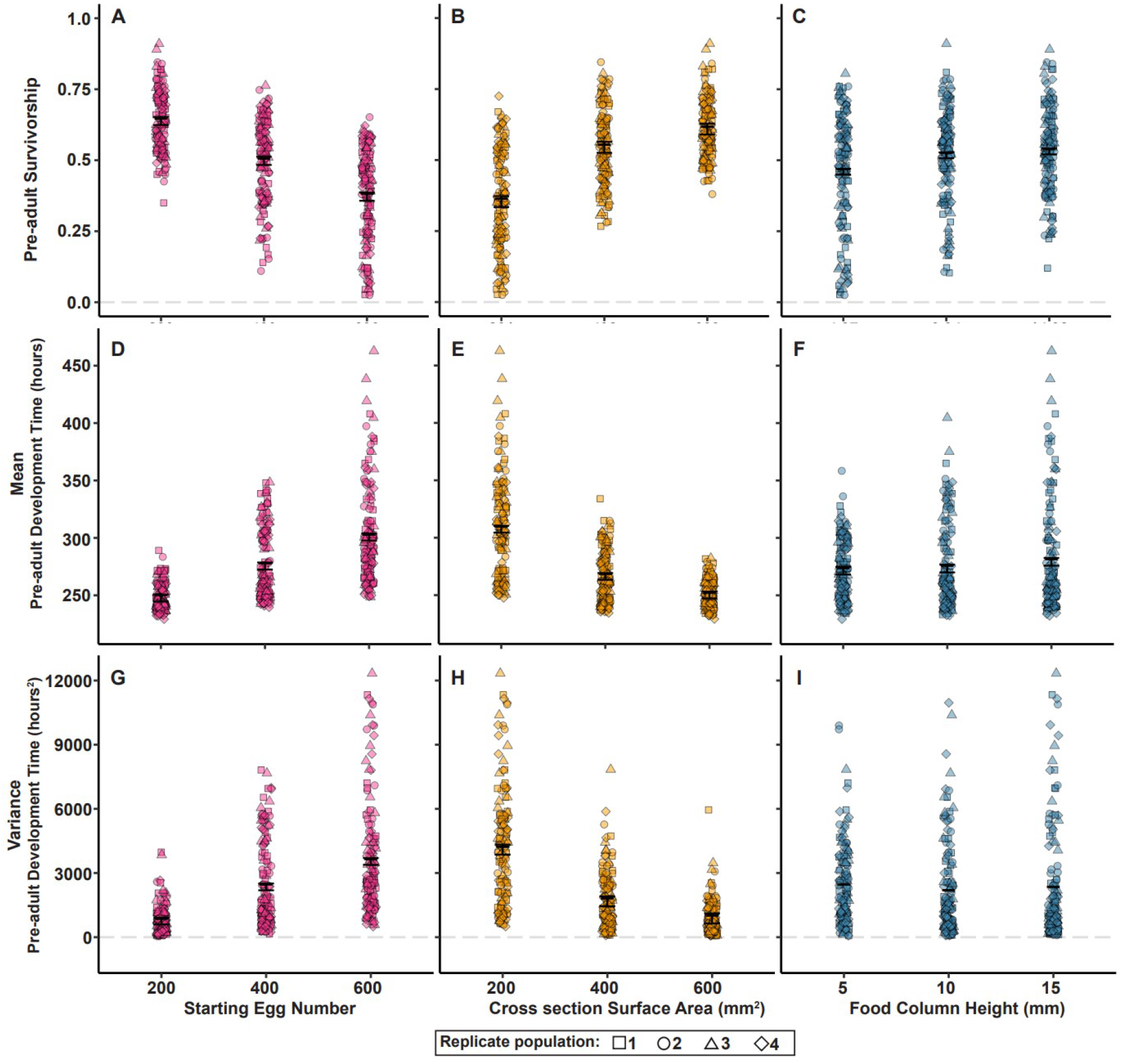
Main effects of the three crowding-inducing factors (egg number, surface area and column height) on mean pre-adult survivorship (A, B, C); mean pre-adult development time (D, E, F); and variance in pre-adult development time (G, H, I), respectively. Error bars represent 95% CI around the means as calculated from Tukey’s post-hoc test. No CI are shown in (I), as there was no significant main effect of column height on variance in development time.

**Figure s2.**
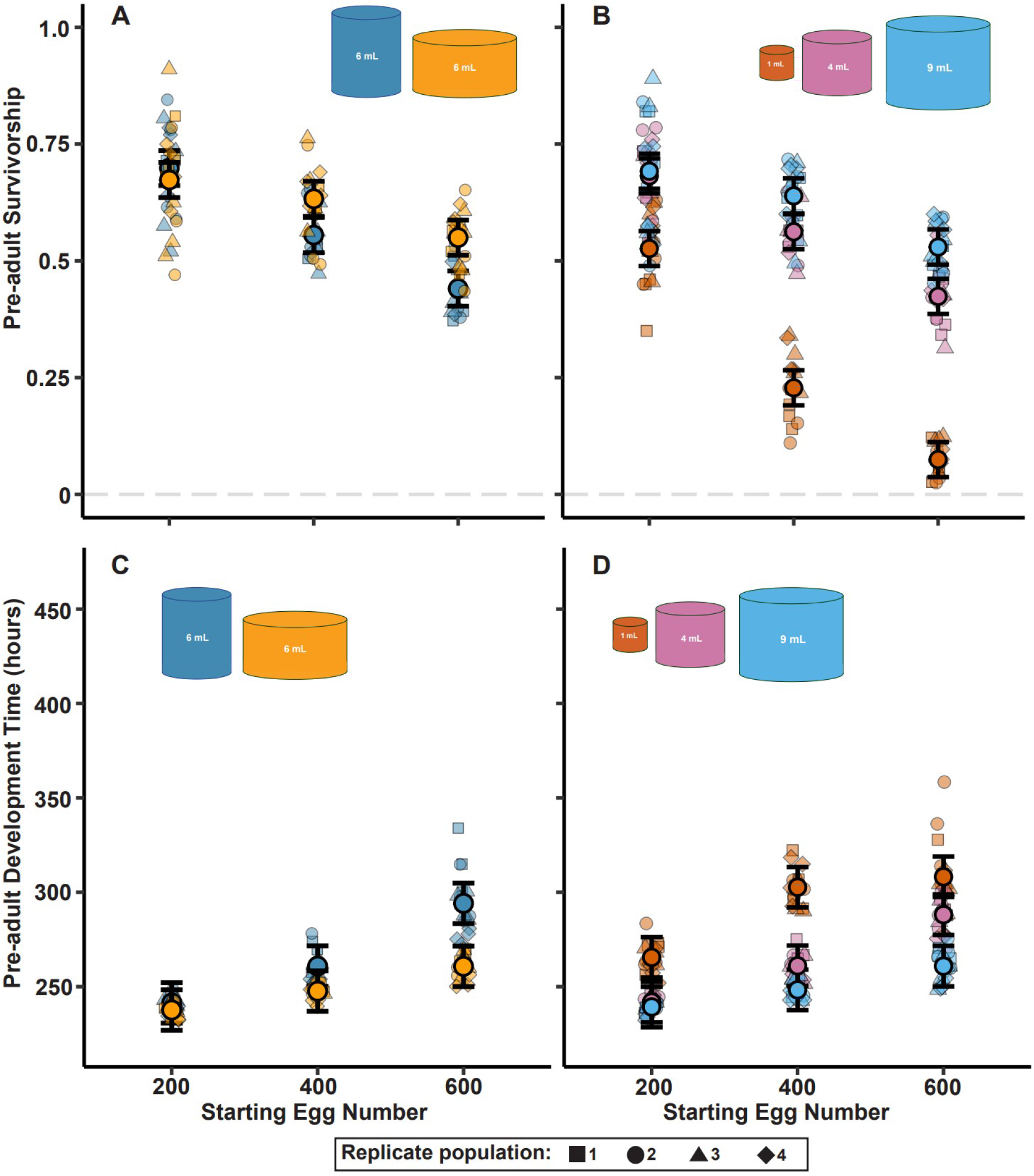
Mean pre-adult survivorship and mean pre-adult development time for (A, C) two variants of the 6 mL cultures; (B, D) Unique cultures of 1, 4 and 9 mL food volumes, for each starting egg number. Error bars represent 95% CI around the means as calculated from Tukey’s post-hoc test.

**Figure s3.**
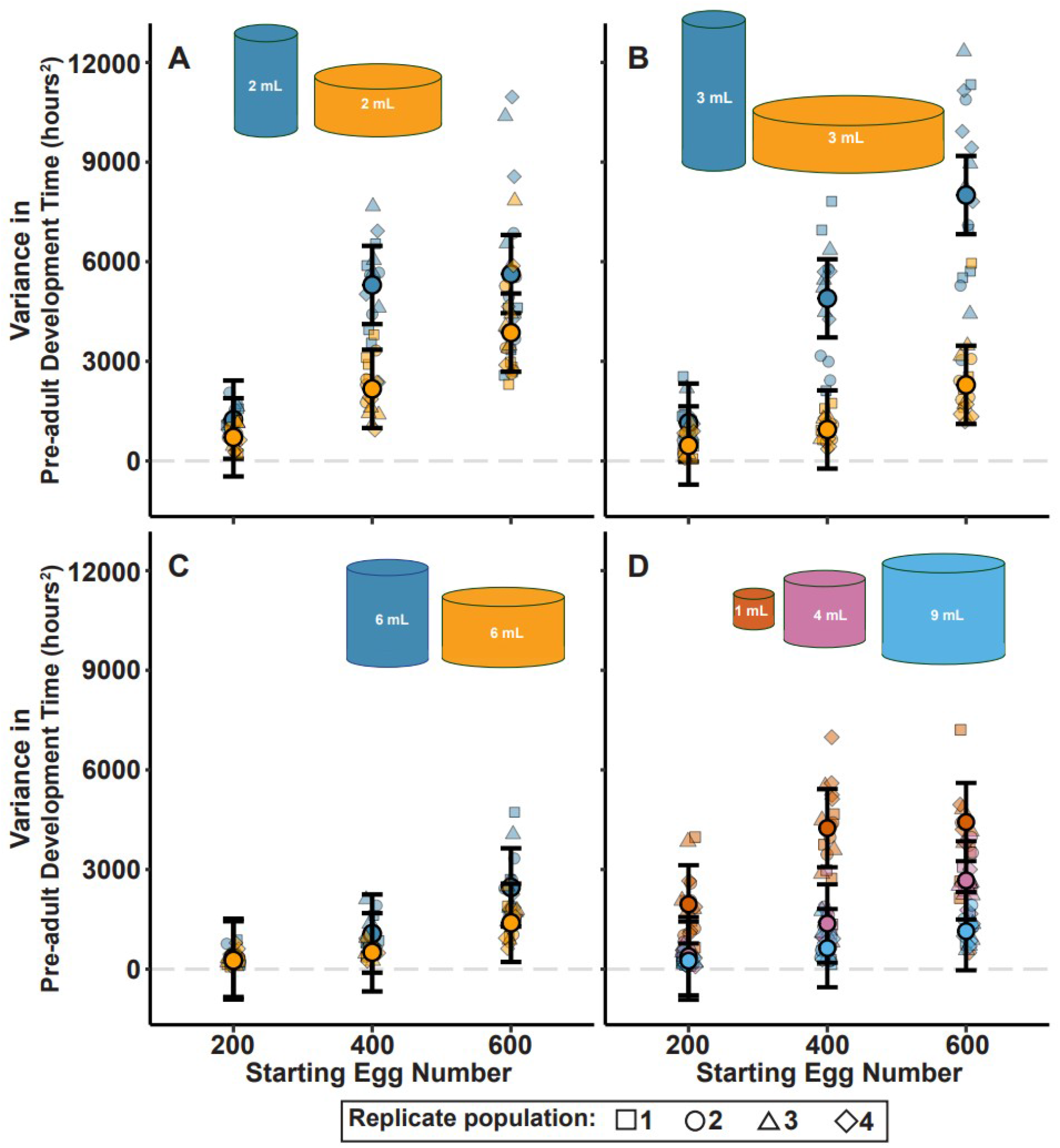
Variance in development time across 200, 400 and 600 eggs seeded in: A. 2 mL food cast in different combinations of surface area and column height; B. 3 mL food cast in different combinations of surface area and column height; C. 6 mL food cast in different combinations of surface area and column height; D. 1, 4 and 9 mL food, respectively. Error bars indicate 95% C.I. around the means, as calculated from Tukey’s post hoc test.

**Figure s4.**
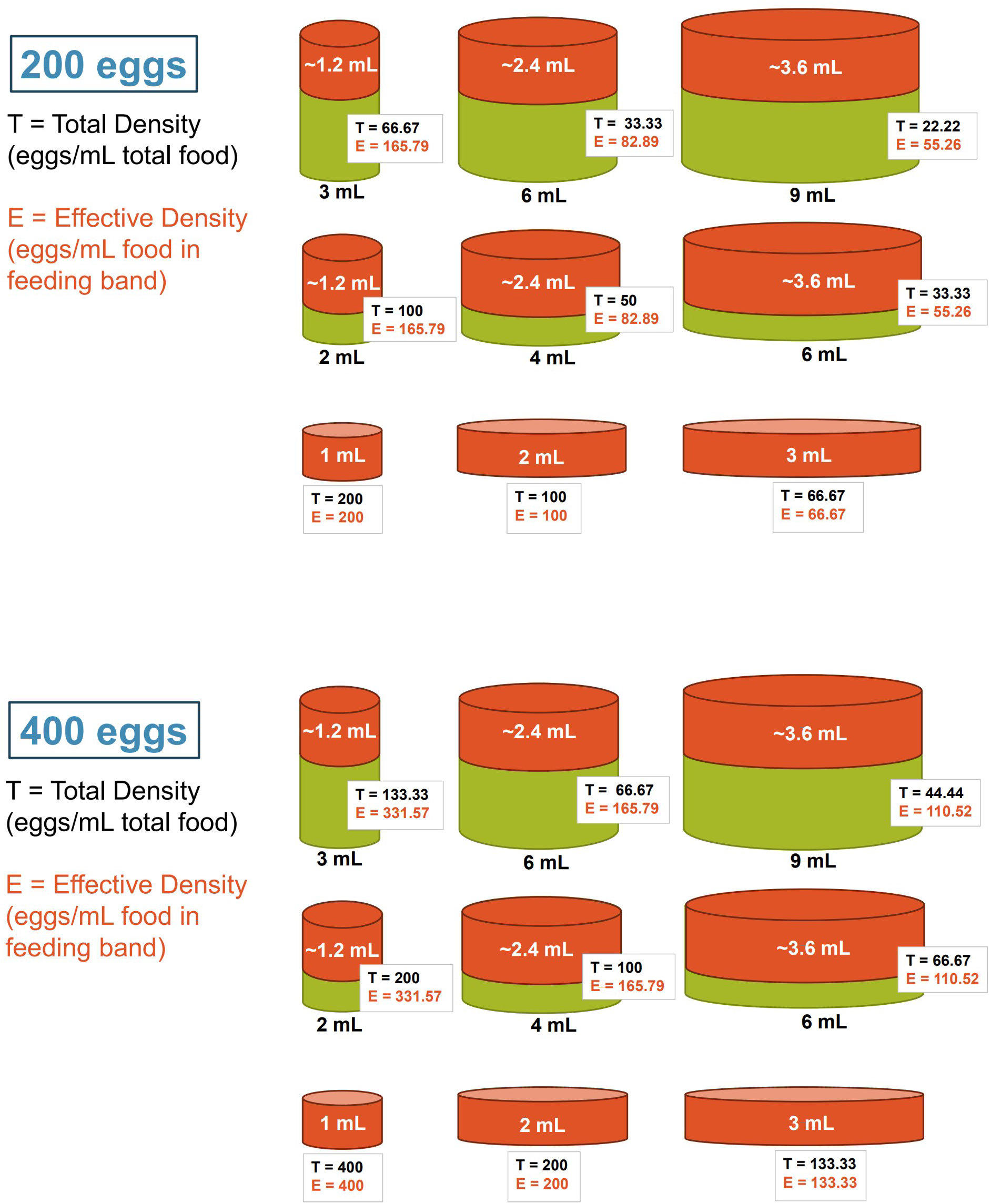
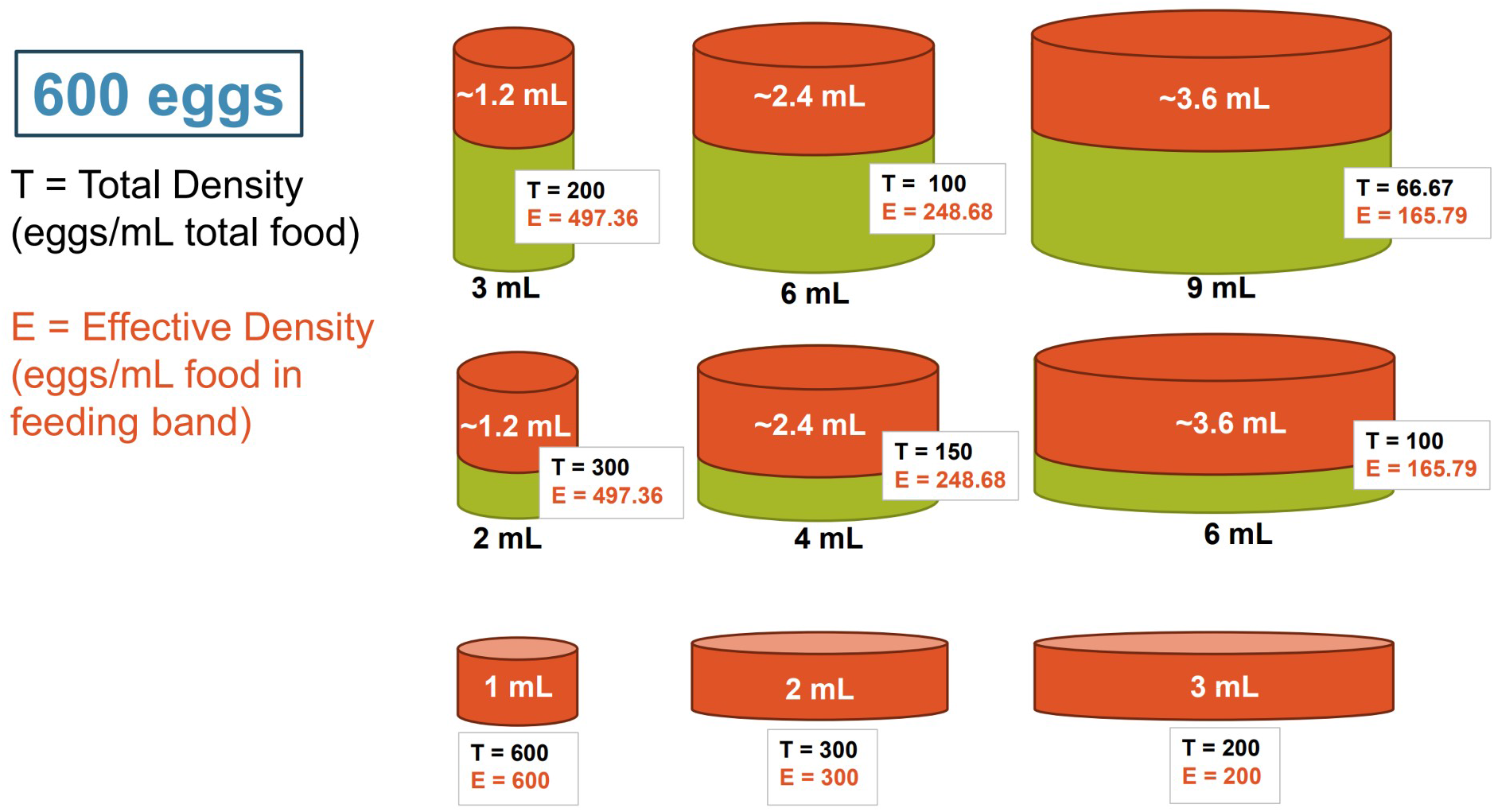
Total density and effective density for each of the 27 treatments used in the experiment.

**Figure s5A.**
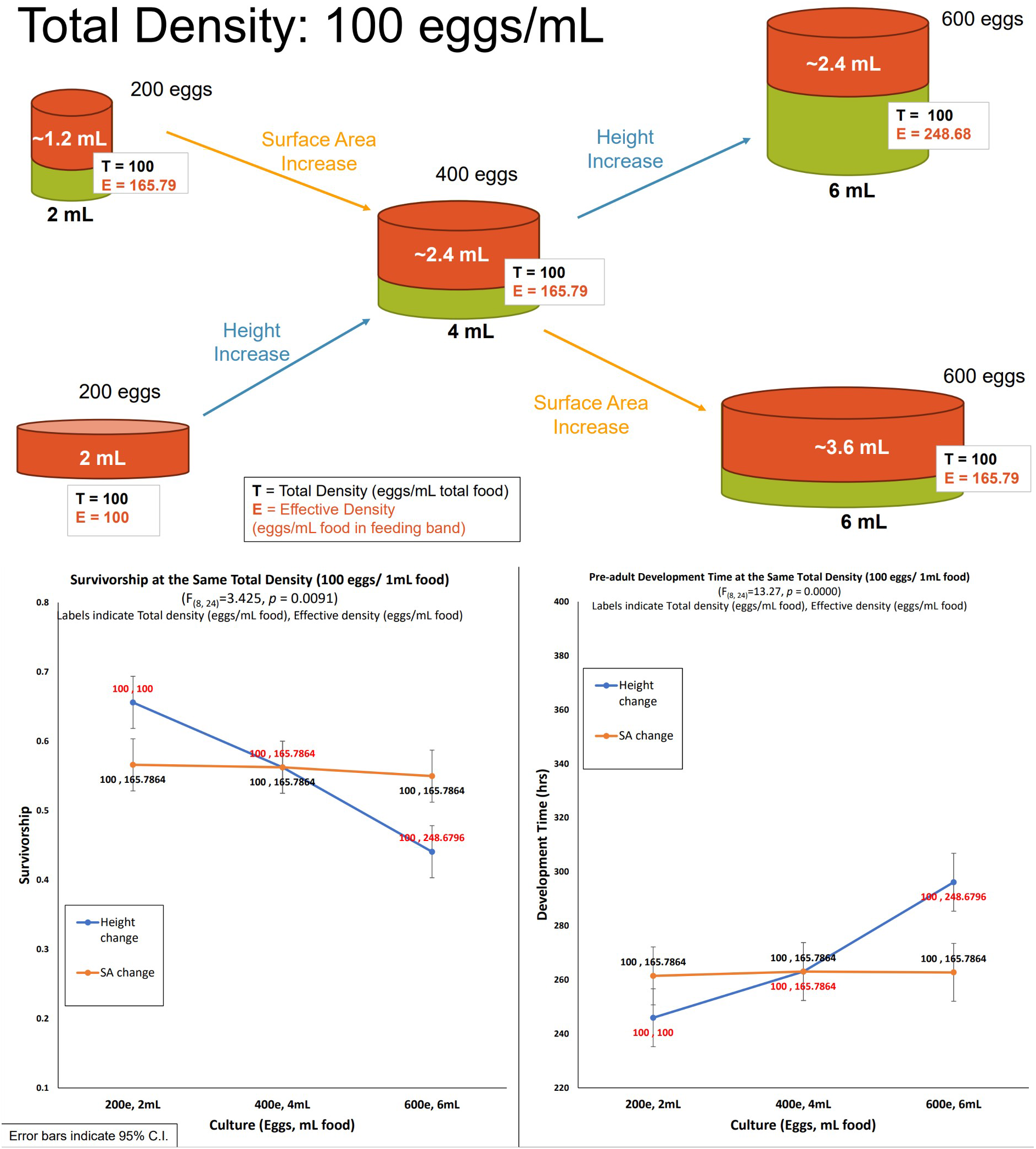
Mean pre-adult survivorship and mean pre-adult development time for each culture with total density of 100 eggs/mL.

**Figure s5B.**
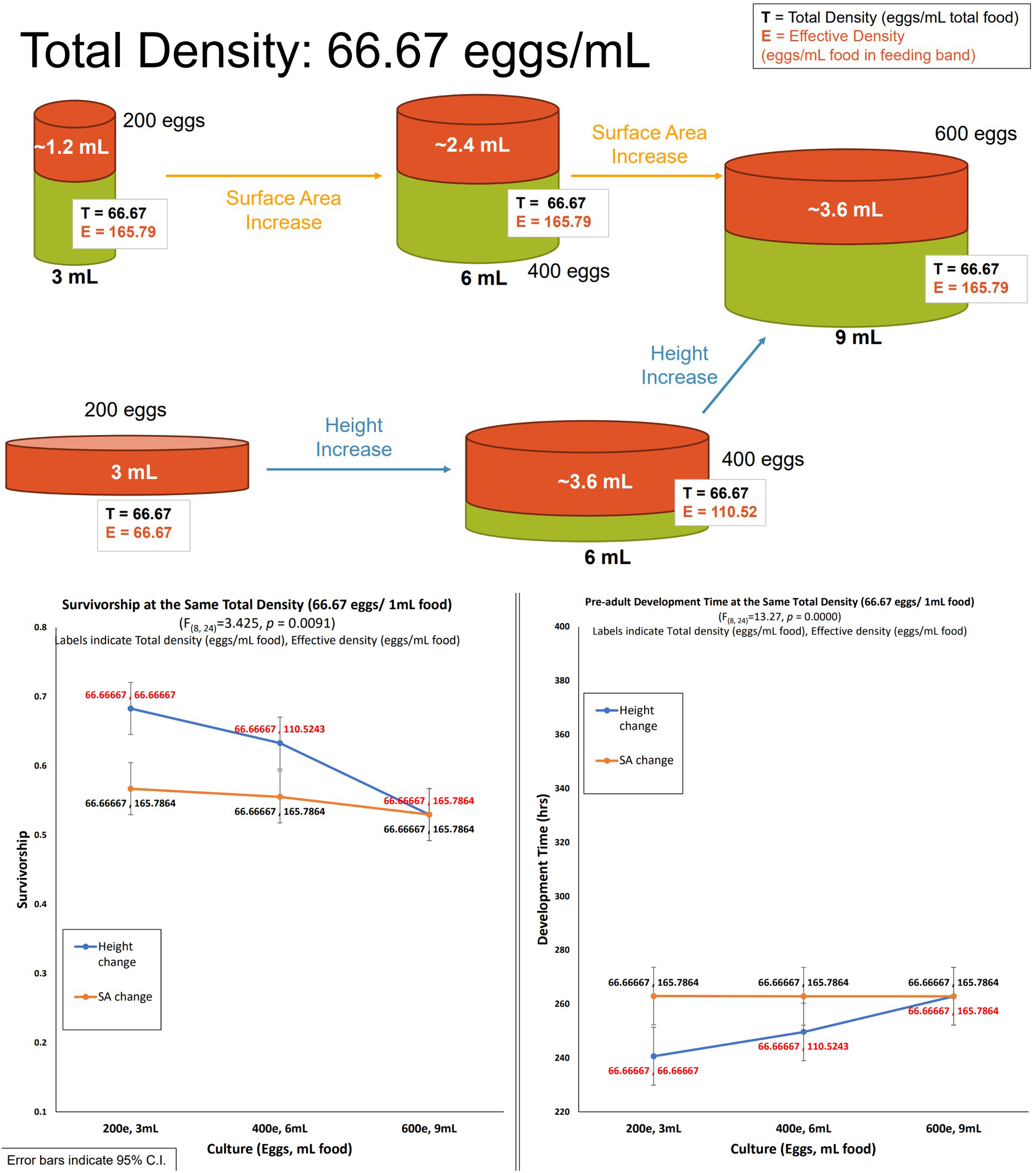
Mean pre-adult survivorship and mean pre-adult development time for each culture with total density of 66.67 eggs/mL.

**Figure 6.**
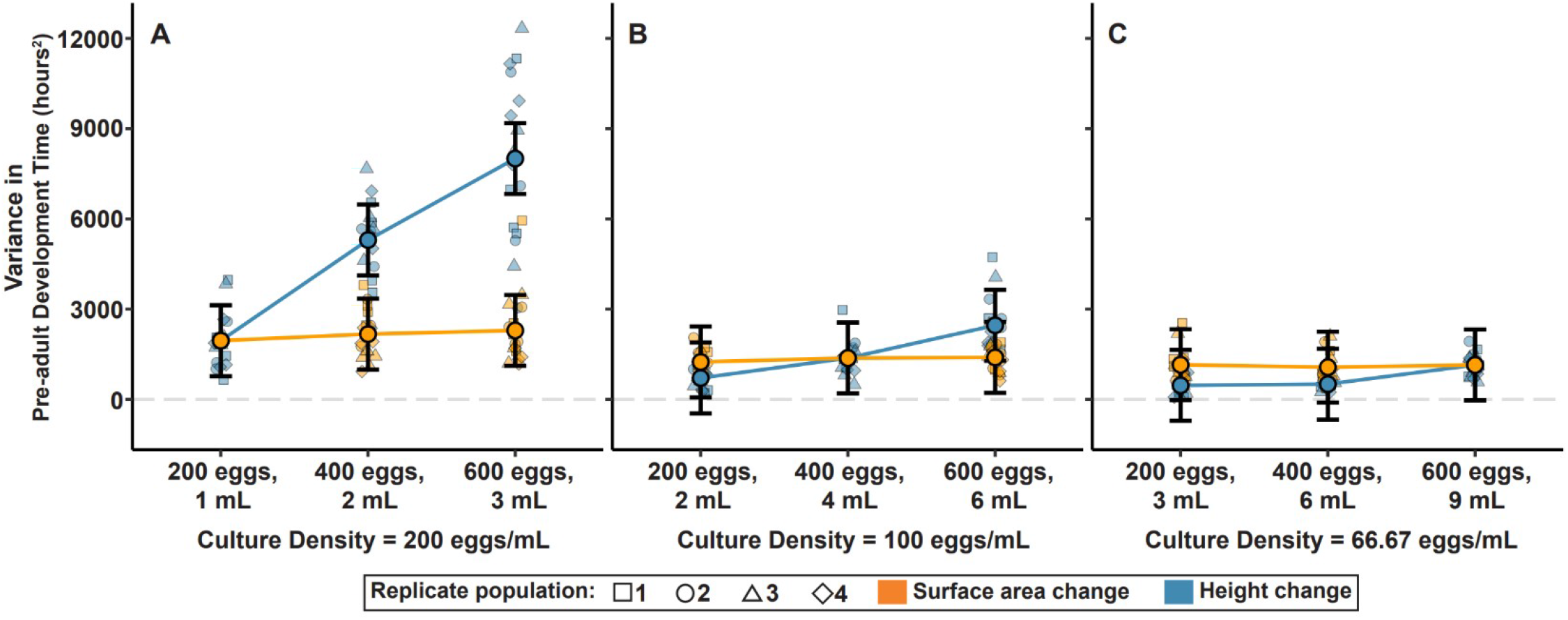
Variance in pre-adult development time for various total density scenarios (see fig. s4): A. Total density of 200 eggs/mL (see fig. 3C); B. Total density of 100 eggs/mL (see fig. s5A);C. Total density of 66.67 eggs/mL (see fig. s5B). Error bars indicate 95% CI around the means, as calculated from Tukey’s post hoc test.

**Figure s7.**
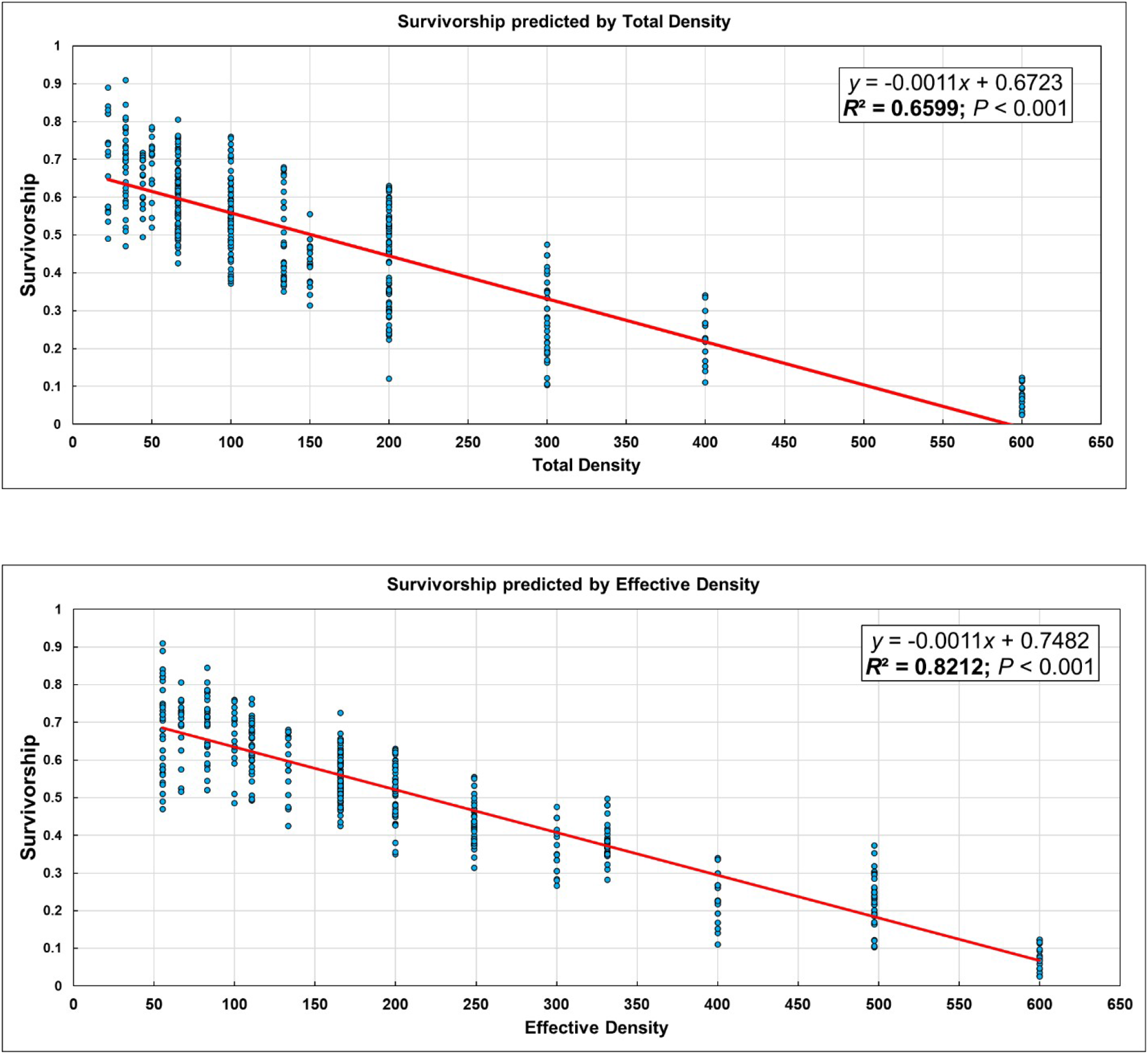
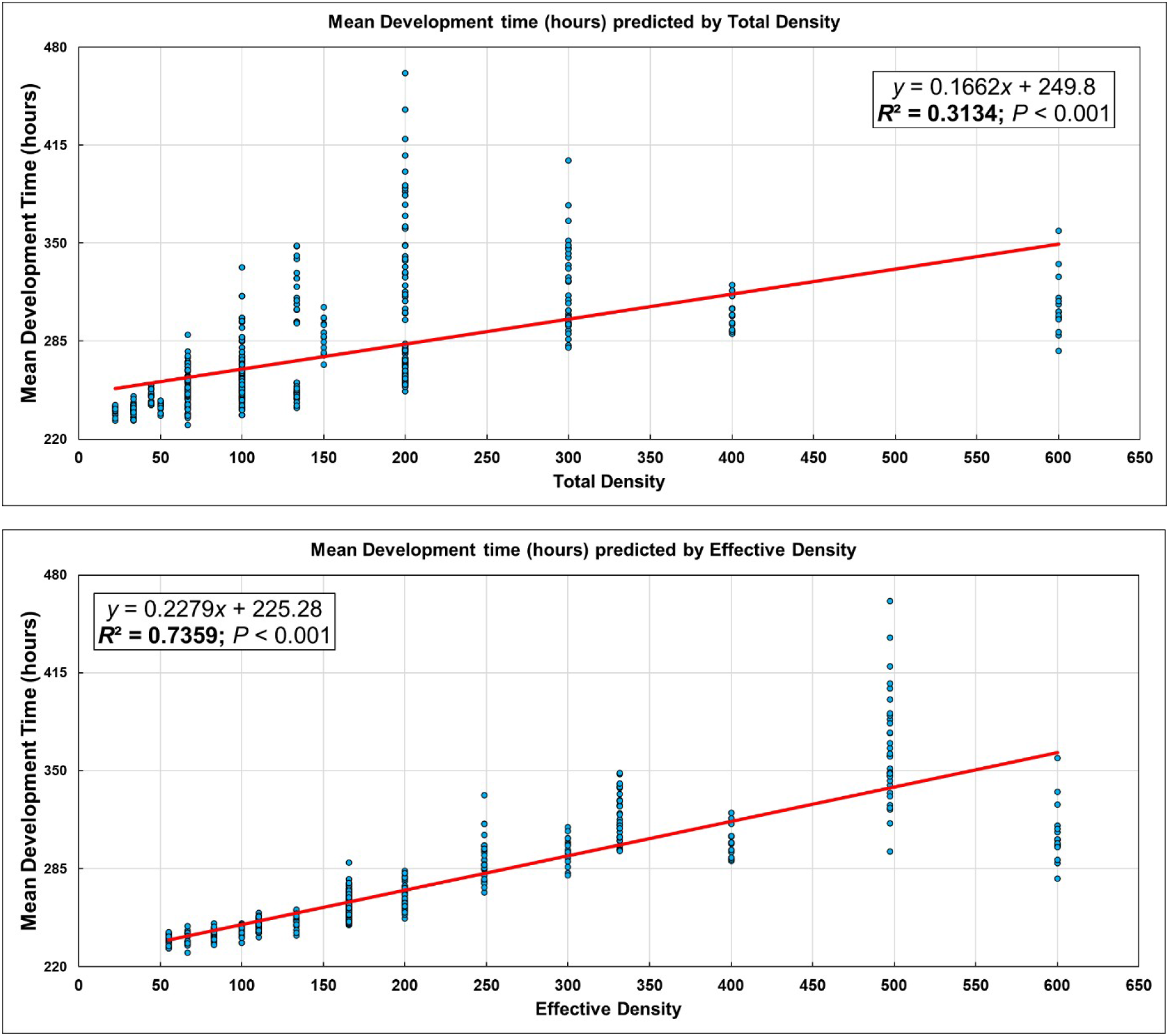
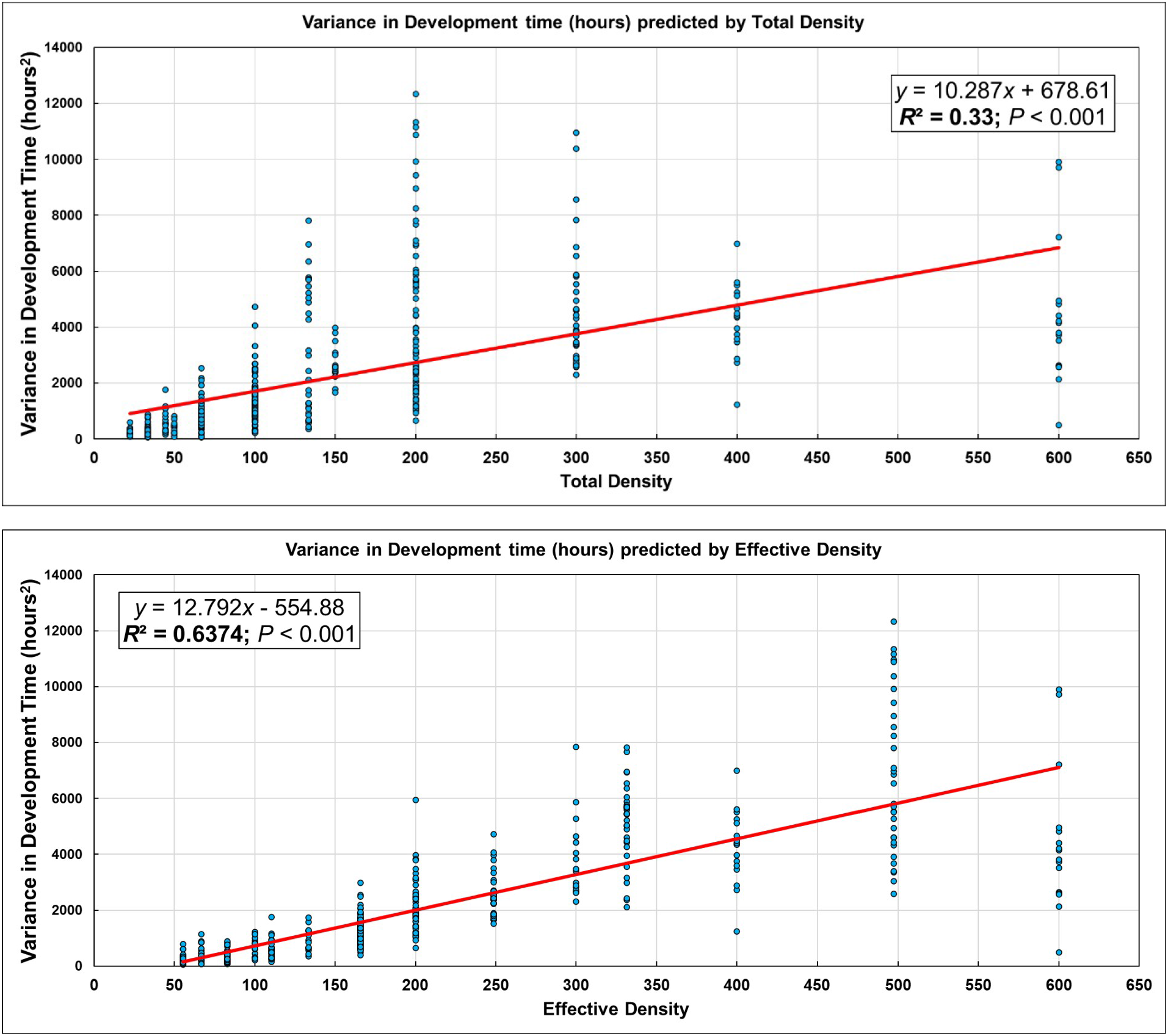
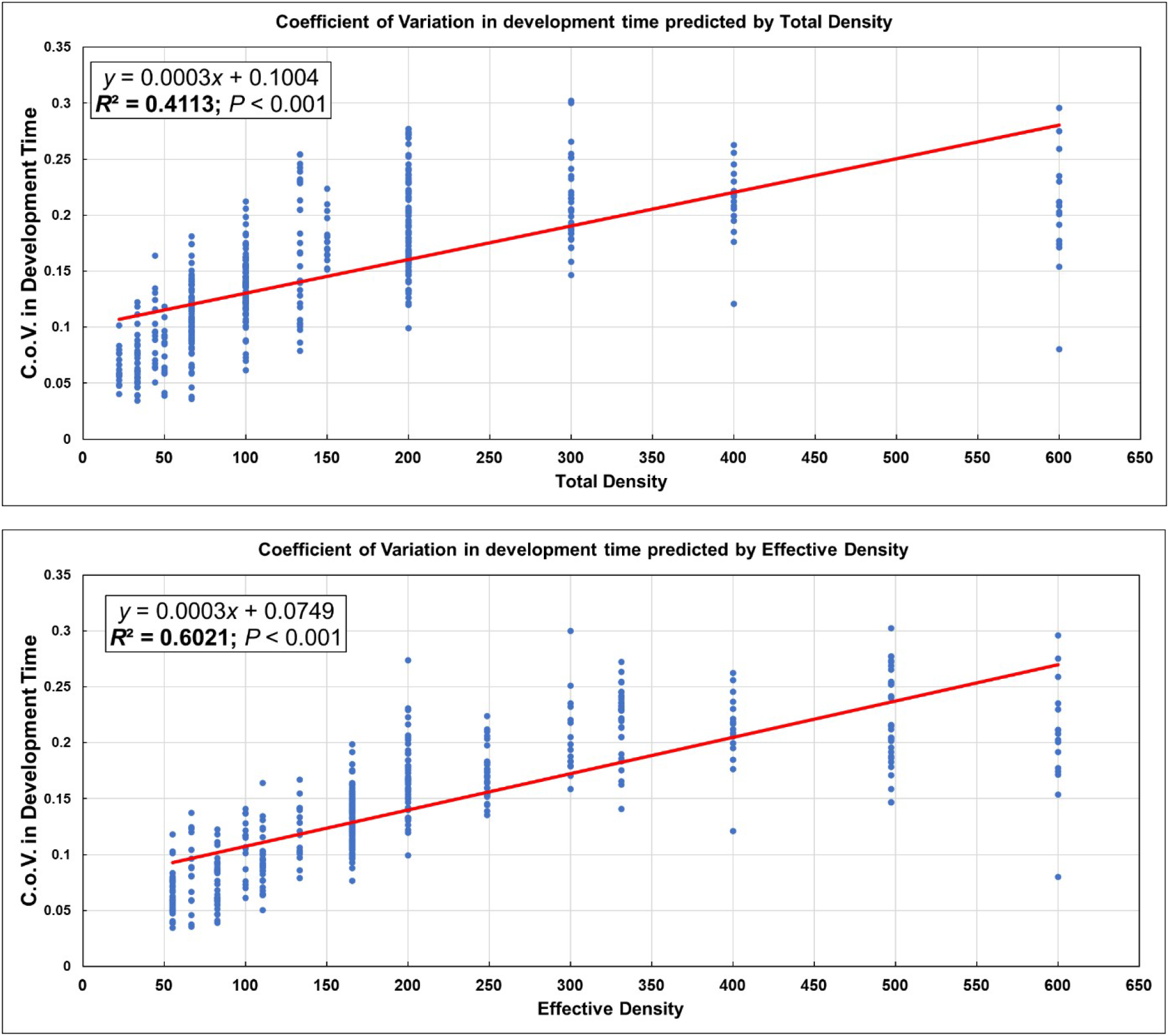
Least-square linear regressions of pre-adult survivorship, mean pre-adult development time, variance of pre-adult development time or coefficient of variation of pre-adult development time as response variables, respectively, as predicted by total density and effective density.

